# The phage stress test: a proving ground for genome language models in biological prediction

**DOI:** 10.64898/2026.09.23.753910

**Authors:** Emily M. Layton, Michael L. Bernauer, Laura D. Weinstock, Eric M. Small, Gina M. Geiselman, George Bachand, Jesse L. Cahill

## Abstract

Reliable biological prediction by AI is most likely to emerge first in the simplest systems with rich datasets and the fastest opportunities for testing and refinement. Phages with small genomes are an ideal testbed. Decades of experimental work have created an unusually information-rich literature benchmark that sits largely outside the sequence repositories typically used to train genome language models (GLMs). Recent work has shown that GLMs such as Evo2 can generate viable whole bacteriophage genomes, demonstrating that genome-scale biological design is possible. The next question is more mechanistic: can such models correctly predict the effects of simple, local sequence changes? Here, we evaluated Evo2 against experimental mutation data from two model phages: the single-stranded RNA phage MS2 and the single-stranded DNA phage ΦX174. We first addressed whether Evo2 sequence log-likelihood scores could distinguish viable from nonviable mutations, and then whether models trained on Evo2 embeddings were predictive of function. Across both phages, Evo2 captured expected sequence-level constraints: stop codons were generally penalized, synonymous substitutions had higher likelihood than nonsynonymous substitutions. Evo2 distinguished among synonymous codons in ways only weakly explained by host codon usage. However, for MS2, these capabilities did not translate into robust biological predictions. Specifically, Evo2 ΔSLL failed to distinguish functional from nonfunctional mutations in the lysis gene, an overlapping viral region that appears to be a particularly hard test case. In ΦX174, performance was stronger but still modest, and much of the apparent signal could be explained by simple covariates such as nonsense mutations, genomic position, and nucleotide distance from the reference. Together, these results introduce phages as a tractable proving ground for stress-testing GLMs against experimentally grounded genotype-to-phenotype tasks. More broadly, they provide a durable framework for identifying what data and model ingredients are required for biologically reliable prediction.

## INTRODUCTION

One of the most compelling visions for biological AI models is its potential to reduce the time, cost, and experimental waste currently required to connect sequence to function [1, 2]. In its strongest form, such a system would help researchers prioritize functional variants, avoid dead-end experiments, and ultimately assist in the design of new biological systems. Yet many investigators remain cautious about committing to available AI toolkits because even a modest rate of wrong predictions can result in net losses for progress once those predictions are tested in the wet lab [3]. The central need, therefore, is for rigorous, information-rich benchmarks that indicate when these tools have matured enough to deliver a true net benefit [4].

Recent advances in genomic language models (GLMs) have shown that models trained on DNA sequence alone can learn biologically meaningful patterns and generate sequences with biological activity [1, 2, 5]. One prominent example is Evo2, a large autoregressive genome foundation model trained on diverse genomic data across all domains of life to learn broad sequence regularities and evolutionary constraints [1]. This work has shown that AI-assisted workflows can generate viable whole bacteriophage genomes in a compact DNA-phage system. In that study, sixteen viable ΦX174-like phages were recovered from 285 synthesized genomes, demonstrating that genome-scale biological design is now possible in principle [5]. Concurrently, it remains unclear how much of that success reflects model-derived biological understanding versus strong human guidance, architecture selection, sequence filtering, and other components of a complex assisted-design workflow. The field has therefore reached an important intermediate stage: AI can participate in successful design pipelines, but it is still not clear whether current models can reliably predict the consequences of simple sequence changes in a way that would let researchers trust them as true biological copilots [2, 6].

Benchmarking AI readiness and credibility is essential for both experimental scientists and policymakers. For researchers, erroneous plausible predictions do not eliminate dead ends; they can simply move those dead ends into more expensive experimental stages [3]. For policymakers, the same question takes on a different form: if AI-enabled genotype-to-phenotype prediction is becoming real, in what biological system would the earliest warning signs appear? We argue that those signals are most likely to emerge first in the simplest tractable biological systems, where the data are deep, experiments are fast, and the cost of being wrong is comparatively low.

An ideal benchmark system for biological AI should satisfy five criteria: (1) it should be intrinsically safe; (2) the genome should be compact enough to support experimental testing and systematic interrogation of sequence variants; (3) it should be supported by a rich body of functional data; (4) it should permit simple experimental readouts such as viability or nonviability; and (5) it should support repeated cycles of prediction, testing, and refinement. Two phages of *E. coli* satisfy these criteria particularly well while probing distinct regimes of biology: the single-stranded RNA phage MS2 and the single-stranded DNA phage ΦX174.

Here, we use published experimental mutation datasets from both systems to benchmark Evo2 against experimentally measured functional outcomes. Many of these datasets already contain viable versus nonviable outcomes for single-nucleotide or single-amino-acid changes [7, 8], but these genotype-phenotype relationships are typically presented in the literature rather than encoded in standard sequence repositories. These extra-corpora data provide a uniquely strong benchmark for asking whether a model can distinguish tolerated from disruptive sequence changes before large new wet-lab training sets are built. Using this framework, we asked what current GLMs capture successfully, where they fail, and what types of data, biological context, and model refinement may be required before predictive success becomes biologically reliable rather than merely computationally plausible.

## RESULTS

### Genome-wide scoring of MS2

To gain a genome-wide picture of Evo2’s scoring of phage mutations, we generated every possible codon variant in MS2, where every codon is changed to each of the 63 other possible codons, and scored each of them with the Evo2 7B model (**Fig. 1)**. Because recent work from our group showed that the current MS2 RefSeq does not represent an experimentally validated infectious genome and identified a minimally corrected, functional sequence [9], we used that corrected sequence as the MS2 reference background for all analyses here (see methods for more detail). As an autoregressive model, Evo2 assigns each nucleotide a probability conditioned on the sequence preceding it, and the sequence log-likelihood is the sum of the logarithms of those probabilities across all positions. Subtracting the reference genome’s score gives a delta sequence log-likelihood (ΔSLL), so a variant with negative ΔSLL is one Evo2 considers less likely than the reference.

**Figure 1.**
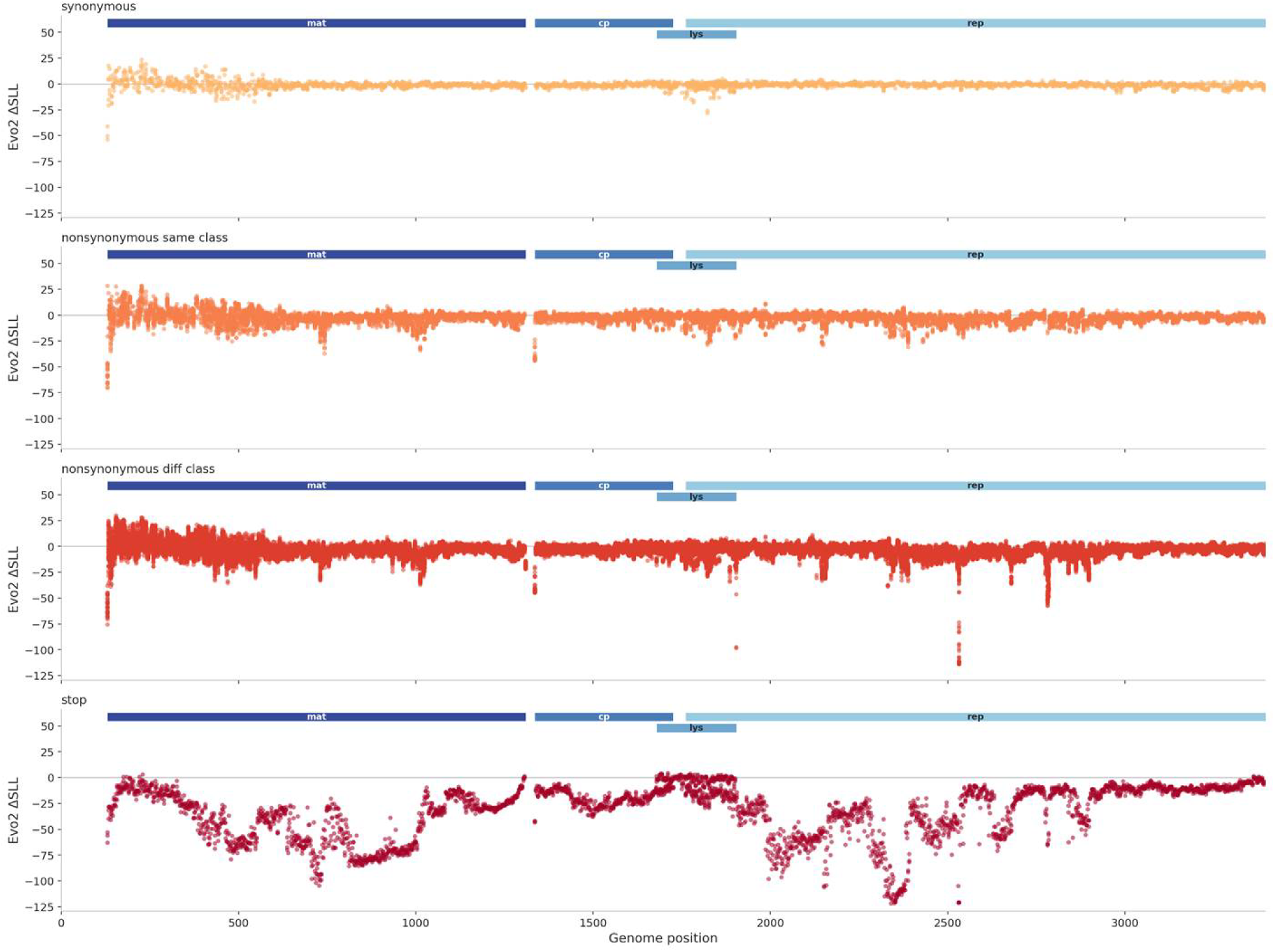
Genome-wide Evo2 scoring of codon substitutions in MS2. For each annotated coding position in the corrected MS2 reference genome, every codon was substituted with each of the 63 alternative codons and scored with Evo2 7B. The y-axis shows ΔSLL, defined as the sequence log-likelihood of the mutant genome minus that of the reference genome; more negative values indicate variants that Evo2 considers less plausible than the reference. The x-axis shows genome position in plus-strand coordinates. Variants are separated into four effect classes: synonymous substitutions, nonsynonymous substitutions that preserve amino-acid class, nonsynonymous substitutions that cross amino-acid class, and nonsense substitutions introducing stop codons. Amino-acid classes were defined as nonpolar, polar, basic, and acidic, with tyrosine and glycine each treated as their own class. Elevated score variability at the 5′ end reflects limited upstream context in autoregressive scoring rather than a biological feature of the genome.

ΔSLL scores separated in generally expected ways: variants with silent mutations clustered around 0 (similar score to the reference), missense mutations were more variable, and nonsense mutations were strongly negative. We observed greater variation among missense mutations that changed the amino acid’s chemical class (polar, nonpolar, basic, or acidic, with tyrosine and glycine each treated separately) than among those that did not (**Fig. 1**). Some of this pattern is expected because synonymous changes are, on average, more similar to the reference sequence than nonsynonymous changes, which may require two or three nucleotide differences. We also noted local hotspots where most changes were scored as especially deleterious, including start codons.

Because Evo2 is autoregressive, each position is scored only in the context of the sequence that precedes it. The 5′ end of the genome is therefore evaluated with minimal context, which produces greater score variability and less constrained ΔSLL values. To verify that the 5’ variation reflected an artifact of autoregressive conditioning rather than a biological prediction, we rescored every sequence with Evo2’s reverse-complement mode, which averages the scores of the forward sequence and its reverse complement (**Fig. S1**). This approach removed much of the 5′ noise but also shifted many scores elsewhere in the genome. Because MS2 is a positive-sense RNA genome, its reverse complement is not the biologically relevant translated or replicated species. We therefore used the reverse-complement analysis only to confirm that the 5′ noise was largely an artifact of limited upstream context and did not pursue those scores further.

These scores generate testable hypotheses about mutations that may be less tolerated; heatmaps showing the ΔSLL score for each codon variant at each site provide higher-resolution views for future investigation (**Fig. S2**). They also reveal substantial variation among synonymous codons at the same site. We therefore asked whether this synonymous-codon variation simply reflected host codon preference. For a phage such as MS2, which depends entirely on *E. coli* translation machinery and must produce high levels of structural proteins within a short infection cycle, host codon usage is a reasonable proxy for one possible source of selective pressure on synonymous sites. To test this, we compared ΔSLL values against *E. coli* codon-usage frequencies within each synonymous codon set. The correlation was positive but weak (ρ = 0.082, p = 4.8 × 10^-6^, n = 3090; **Table 1**) and was consistent in direction across all four MS2 genes. However, it accounted for less than 1% of the rank variance in the ordering of synonymous codons by ΔSLL, meaning that Evo2’s codon preferences were only minimally explained by host usage. Thus, Evo2 distinguishes among synonymous codons largely along an axis unrelated to simple *E. coli* codon bias.

**Table 1.** Evo2 distinguishes among synonymous codons in MS2, but *E. coli* codon usage explains little of that separation. For each synonymous codon, we compared its Evo2 score (ΔSLL) with the frequency with which that codon is used in *E. coli* relative to its synonymous alternatives. Positive Spearman correlations indicate that codons more frequently used by *E. coli* tend to receive higher Evo2 likelihood scores.

| GENE | COUNT | SPEARMAN CORR. | P-VAL |
| --- | --- | --- | --- |
| CP | 333 | 0.102 | 0.0640 |
| LYS | 219 | 0.156 | 0.0209 |
| MAT | 1070 | 0.067 | 0.0282 |
| REP | 1468 | 0.079 | 0.0023 |
| ALL GENES | 3090 | 0.082 | 4.79E-06 |

### The challenge of the MS2 lysis gene

To investigate whether Evo2 captures complex sequence constraints (e.g., overlapping coding regions), we examined its scoring of the MS2 lysis gene, which occupies a compact, high-information-density region of the genome, with three converging reading frames. Its coding sequence overlaps other functional sequence space, with its N-terminus embedded in the coat gene and its C-terminus embedded in the replicase. Interposed within this same genomic neighborhood is the MS2 operator, a defined RNA hairpin bound by coat protein. The operator contributes to at least two critical functions: it helps nucleate coat-protein assembly and regulates access to replicase translation [10]. As a result, this region simultaneously encodes protein sequence, preserves overlapping reading frames, and maintains RNA structures required for gene regulation. It is therefore one of the densest and most functionally layered regions in the MS2 genome, where disruption of any one component is likely to produce a clear, often lethal, phenotype.

We noticed that variants introducing stop codons in the lysis gene received ΔSLL values conspicuously near zero, in contrast with other MS2 genes where stop codons were generally assigned much more negative ΔSLL values (**Fig. 1, Fig. S2C, Fig. S3, Fig. S4**). One explanation is that Evo2 is responding more strongly to the overlapping coat and replicase constraints than to lysis-specific loss of function. In many cases, a mutation that introduces a stop codon in the lysis frame produces only a nonsynonymous substitution in the overlapping coat or replicase context. If Evo2 has seen many viral sequences encoding related coat proteins and replicases, but relatively fewer comparable lysis proteins, it may weigh the former more strongly than the latter. This is plausible because lysis proteins are often short, highly divergent, and functionally specialized [11].

Prior work showed that when the lysis protein is studied in isolation from the full genome, the N terminus is dispensable whereas the C terminus is essential [12]. As a result, truncating mutations would be expected to be strongly deleterious whenever they prevent translation of the essential C-terminal region, with only the most distal stop codons plausibly retaining partial function. That expectation was not reflected in Evo2 scoring. A dispensable N terminus might be expected to tolerate a broader range of substitutions than the essential C terminus, but that pattern was not observed (**Fig. S2C**). Although this does not by itself prove that Evo2 lacks useful information about the lysis gene, it does suggest that the model is not recognizing the functional constraints of *L* in the way one would expect.

### ΔSLL does not predict lysis protein functionality

To test whether Evo2 ΔSLL scores correspond with experimentally measured function, we next compared them with published mutation data for the MS2 lysis gene. Chamakura et al. [8] performed random mutagenesis of the 75-aa lysis protein L and identified single nucleotide mutations that either preserved or abolished lytic function. This dataset provides a useful benchmark because it links defined mutations to experimentally measured functional outcomes, allowing a direct test of whether Evo2 model-derived sequence scores track biological effect. In that study, mutant L alleles were expressed from an inducible plasmid and surviving colonies were initially selected as candidates for loss of lysis. Because functional L kills the host, survivors were presumed to be lysis-defective; however, follow-up testing in liquid culture showed that a subset of these mutants still lysed normally, allowing the dataset to distinguish truly nonfunctional variants from variants that appeared to support lysis in the original selection.

We hypothesized that mutations abolishing L function would receive lower ΔSLL values than mutations retaining function. This was not the case; ΔSLL did not significantly differ between functional and nonfunctional variants (p=0.79, AUROC=0.44, n=83; **Fig. 2A**); where the normalized U statistic (i.e., AUROC) was used to allow comparisons across tests given the sensitivity of the Mann-Whitney U statistic to sample size. The most negative ΔSLL values in the nonfunctional group were all nonsense mutations, but even after separating nonsense and missense variants, neither subgroup differed from the functional variants (nonsense: p-adj=0.79; missense: p-adj=0.71; **Fig 2A**). ΔSLL also showed no relationship to measured L protein production in the subset of variants with available western blot data [8] (p=0.85, AUROC=0.48, n=60; **Fig. 2B**).

**Figure 2.**
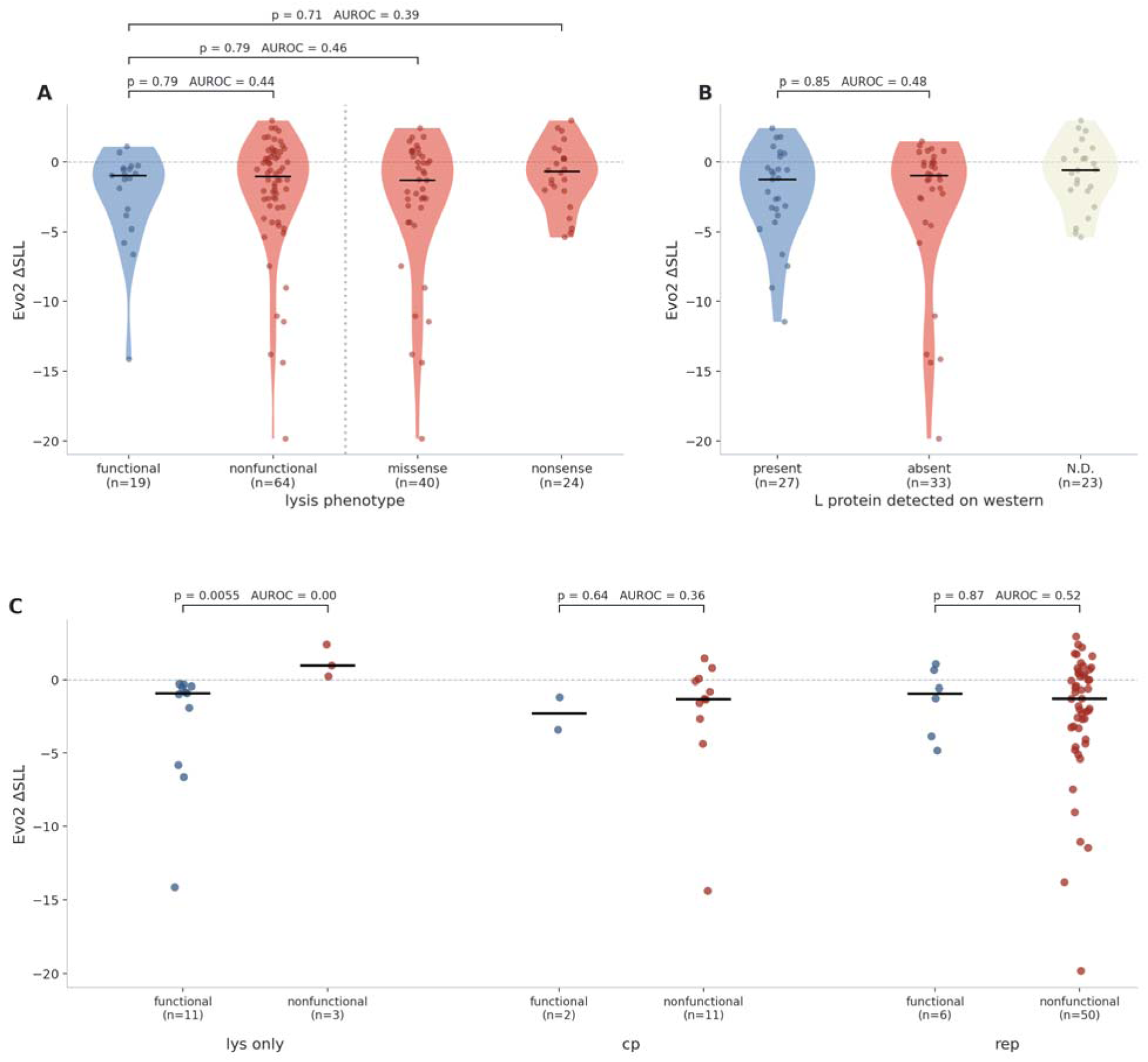
Evo2 ΔSLL does not distinguish lysis-competent from lysis-defective MS2 variants. **(A)** ΔSLL for 83 distinct single-nucleotide variants in the MS2 lysis gene, grouped by lysis phenotype from Chamakura et al. (2017). To the right of the dotted divider, the nonfunctional class is further subdivided by whether the mutation introduces a stop codon. Violin plots show the distribution of scores in each group, black bars indicate medians, and points represent individual variants. Brackets show two-sided Mann–Whitney comparisons of each group against the functional class, with Holm correction across the three comparisons. **(B)** The same variants grouped by whether L protein was detected by western blot. **(C)** The phenotype comparison from panel A repeated within each reading frame. Because L overlaps cp and rep, most nucleotide substitutions affect more than one coding context. Points and median bars are shown without violin plots because some groups contain only two variants. Brackets show within-frame comparisons and are uncorrected across the three frames. AUROC is oriented so that values above 0.5 indicate the first group listed in each comparison scores higher on average; values below 0.5 indicate the opposite. In panels A and C, the first group is the lysis-competent class. In panel B, the first group is variants with detectable L protein.

Because scores for L mutations may be influenced by their effects on the overlapping coat and replicase reading frames, we next stratified variants by whether the mutated position lay only within L or also overlapped coat or replicase. Surprisingly, overlap did not fully explain the mis-scoring. Among the 14 variants within the non-overlapping region of L, ΔSLL was significantly lower for viable variants than for non-viable variants (Mann–Whitney U=0, p=0.0055, AUROC=0.00, n=12; **Fig 2C**). Together, the significant group separation and AUROC of 0.00 indicate a reversed ranking, with non-viable variants receiving higher ΔSLL scores than viable variants. Although this subset is small and requires cautious interpretation, it shows that Evo2’s difficulty with L cannot be attributed solely to overlap with neighboring genes. Even at positions unique to the lysis reading frame, Evo2’s ranking remained inconsistent with the experimental phenotype.

### Models of MS2 intermediate embeddings

Because ΔSLL did not predict MS2 lysis outcomes, we next assessed whether Evo2’s internal sequence representations (embeddings and intermediate layer space), carried information about phenotype that was not accessible through sequence likelihood alone. From each of the 32 processing blocks of the Evo2 7B model, we extracted intermediate embeddings under three pooling schemes: the coding sequence, the mutated site, and the whole genome (32 blocks × 3 pooling schemes = 96 configurations; **Fig. 3A**). We then trained a logistic regression model on these embeddings using five-fold grouped cross-validation using functional vs non-functional designations from Chamakura et al. as the target variable. Variants were grouped by site such that different substitutions at the same nucleotide position were always placed in the same fold. This prevented the model from training on one substitution at a site and then being tested on another substitution at that same site.

**Figure 3.**
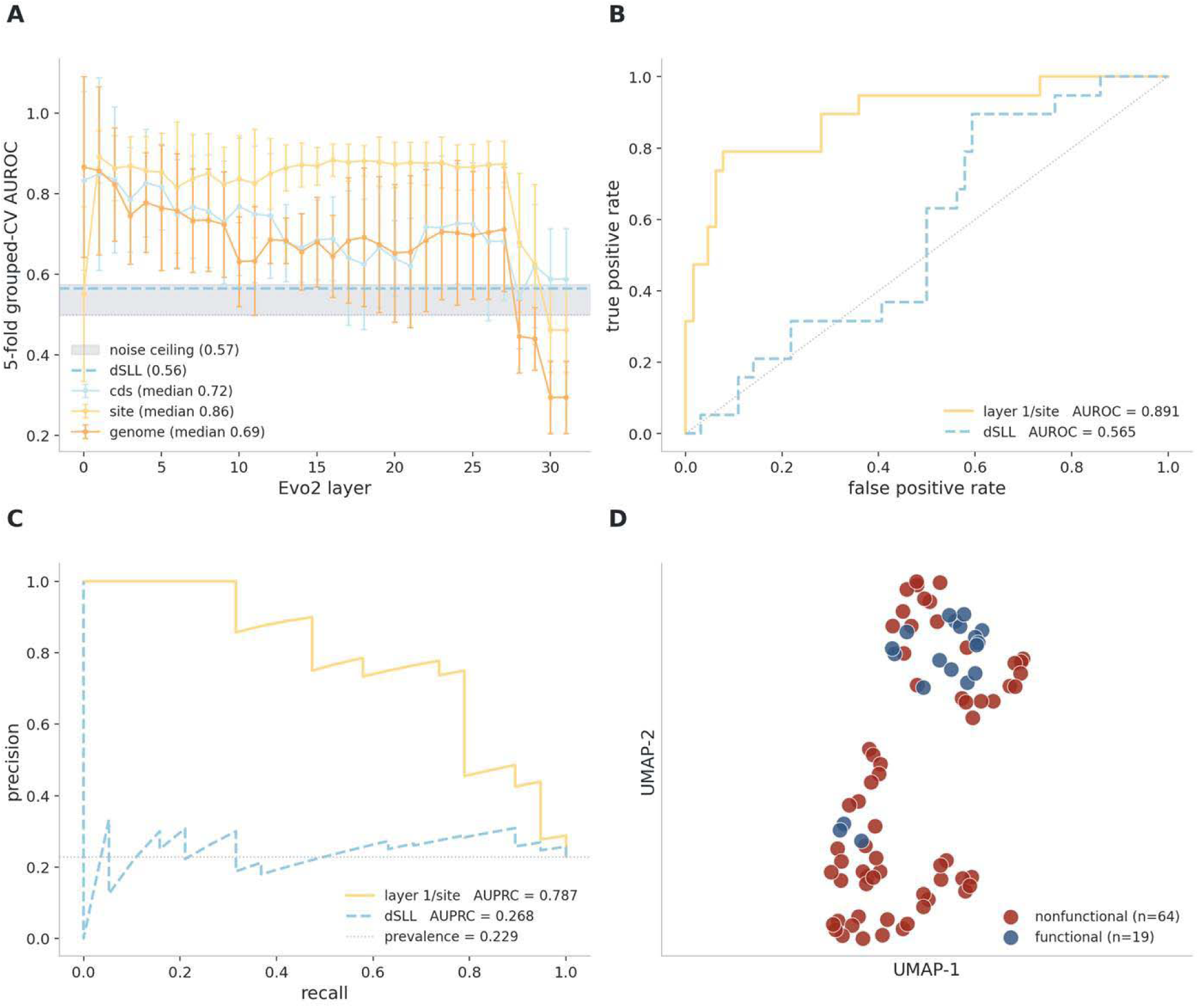
Intermediate Evo2 embeddings improve MS2 lysis classification relative to ΔSLL, but do not cleanly separate phenotype. (A) AUROC from grouped five-fold cross-validation across Evo2 layers for the three pooling schemes tested: coding sequence (CDS), mutated site, and whole genome. Blue dashed line represents ΔSLL baseline. Though ΔSLL ranks the two classes backwards, as reported in Fig. 2A (AUROC = 0.435), here it is reported in its better-performing orientation (1 – AUROC = 0.565) to allow fair comparison with orientation-agnostic fitted models. Grouping was performed by mutation site so that different substitutions at the same position were always kept in the same held-out subset. (B) Receiver operating characteristic (ROC) curve for the best-performing model configuration, layer 1 with site pooling (yellow), compared with ΔSLL used directly as a score (blue dashed). (C) Precision-recall curve for the same comparison as in panel B. The gray dotted line indicates class prevalence. (D) UMAP projection of the best-performing embedding representation (layer 1, site pooling). Each point represents a variant; blue points indicate functional variants and red points indicate nonfunctional variants. Although structure is visible in the embedding space, the projection does not cleanly separate the two phenotype classes.

Performance was broadly similar across most model configurations. Specifically, cross-validation variability was comparable to the performance differences seen across most processing blocks and pooling schemes. Nevertheless, site pooling consistently outperformed the other two schemes (coding sequence, and whole genome) across much of the model, with the best logistic regression results derived from block 1 (AUROC = 0.891 ± 0.065; **Fig 3A**). For that configuration, both the receiver operating characteristic curve (ROC) and precision-recall curves suggested that a model trained on embeddings captured more information about phenotype than ΔSLL alone (**Fig. 3B,C**). UMAP visualization of the same embeddings produced two visible clusters in the first two dimensions, but these did not correspond cleanly to functional and nonfunctional classes (**Fig. 3D**). To test whether the embeddings contained phenotype signal that was not accessible to a simple linear classifier, we also evaluated three more flexible alternatives on the same best-performing representation: a random forest, which can capture rule-like interactions among features; a support vector machine with a radial basis function kernel, which can fit curved decision boundaries; and a multilayer perceptron, a small neural network capable of learning more flexible nonlinear mappings. None substantially improved upon the logistic regression model (**Table 2**).

**Table 2.** Classifier comparison on MS2 embeddings (block 1, site pooling). Grouped five-fold cross-validation; 19 functional and 64 nonfunctional variants across 49 sites. “SD” is the standard deviation across folds. ΔSLL is a training-free baseline. Though ΔSLL ranks the two classes backwards, as reported in **Fig. 2A** (AUROC = 0.435), here it is reported in its better-performing orientation (1 – AUROC = 0.565) to allow fair comparison with fitted models, which can learn either score orientation.

| CLASSIFIER | AUROC | AUPRC | SD |
| --- | --- | --- | --- |
| ΔSLL | 0.565 | 0.268 | |
| LOGISTIC | 0.891 | 0.787 | 0.064 |
| RANDOM_FOREST | 0.826 | 0.668 | 0.136 |
| SVM_RBF | 0.825 | 0.704 | 0.108 |
| MLP | 0.9 | 0.791 | 0.101 |

### The MS2 Model result may be explained by genomic position

Although we attempted to mitigate information leakage into the training/test sets by grouping cross-validation folds on mutation site, the embeddings contain positional information which may be contributing to the model’s performance. To test whether the model result reflected true phenotype signal or simpler properties such as SNP position, we refit the same classifiers using only basic covariates (that is, simple non-embedding features of each variant, such as genomic position and whether the mutation introduces a stop codon) under the same grouped cross-validation scheme used for the models (**Table 3**). Using this procedure, all mutations at the same site were kept together in the same held-out subset, so the classifier was never trained on one substitution at a site and then evaluated on another substitution at that same site. Under a random forest, genomic position alone reached AUROC 0.905, and position combined with stop status reached AUROC 0.970. Adding embeddings to these covariates did not recover additional signal (AUROC 0.843). This interpretation should be made cautiously, however, because the phenotype labels derive from an isolated-gene lysis assay rather than from whole-genome infectivity. If the embeddings capture broader overlapping or genome-context constraints, then some apparent misclassifications may reflect a mismatch between the assay label and whole-genome biological effect rather than a simple failure of the model. We therefore do not take the MS2 model results as clear evidence that Evo2 has learned the functional logic of the lysis gene. Rather, much of the apparent model performance can be explained by where a mutation occurs in the gene, rather than by the model capturing the biological effect of the mutation itself.

**Table 3.**
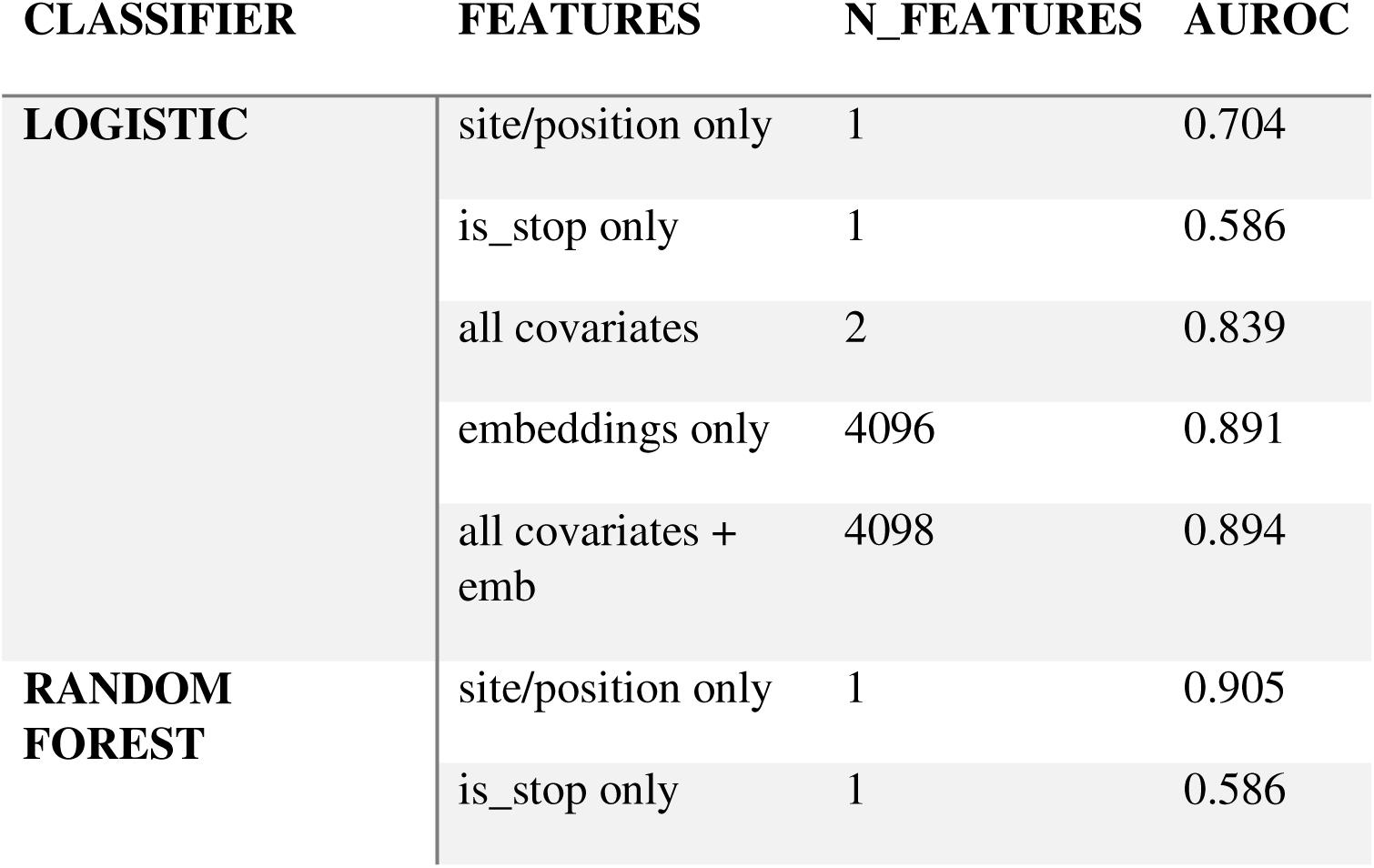
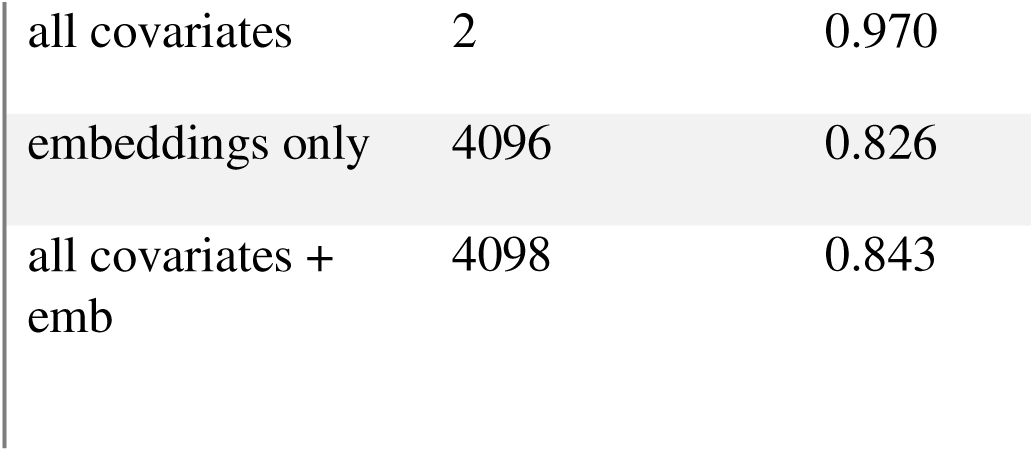
Simple covariates explain much of the apparent MS2 embedding signal. Each row fits a classifier on a specific feature set under the grouped five-fold cross-validation scheme used for the embedding models, so rows differ only in the information provided to the classifier. “Covariates” are the genomic position of the substitution and whether the mutation introduces a stop codon; “embeddings” are the 4096-dimensional vectors of the best-performing Evo2 representation (block 1, site pooling). AUROC is oriented so that values above 0.5 indicate that functional variants tend to receive higher scores than nonfunctional variants. A feature set is interpreted as contributing information beyond another only if the combined model outperforms each input considered separately.

| CLASSIFIER | FEATURES | N_FEATURES | AUROC |
| --- | --- | --- | --- |
| LOGISTIC | site/position only | 1 | 0.704 |
|  | is_stop only | 1 | 0.586 |
|  | all covariates | 2 | 0.839 |
|  | embeddings only | 4096 | 0.891 |
|  | all covariates + emb | 4098 | 0.894 |
| RANDOM FOREST | site/position only | 1 | 0.905 |
|  | is_stop only | 1 | 0.586 |

|  |  |  |
| --- | --- | --- |
| all covariates | 2 | 0.970 |
| embeddings only | 4096 | 0.826 |
| all covariates +<br>emb | 4098 | 0.843 |

One possible reason Evo2 struggles in this setting may be due to limited representation of similar sequences in training. To explore this, we built a local BLAST database from the OpenGenome2 dataset used to train Evo2 and searched for sequences similar to the MS2 reference. This yielded only nine related sequences (**Supplementary Table 1**), consistent with the idea that MS2-like lysis sequences may be sparsely represented in the training corpus.

Taken together, these results suggest that the MS2 lysis region is not merely a difficult edge case, but a useful stress test for genome language models. It compresses a short, functionally specialized protein, overlapping coding relationships, and RNA regulatory features into a very short sequence, making it the kind of high-information-density region in which a sequence can appear plausible to Evo2 while still being functionally wrong in biology.

### Genome-wide scoring of ΦX174

Evo2 is a DNA-centric GLM [1] and may therefore be less well suited to phenotype prediction in systems where RNA-level constraints (e.g., RNA folding) could dominate biological outcome. To address whether the performance seen in MS2, an RNA phage, is a result of model mismatch and genome type, we repeated the analysis on ΦX174, a single-stranded DNA phage that infects *E. coli*. ΦX174 has a similar sized genome to MS2, providing a complementary benchmark that may be more closely aligned with Evo2’s DNA training corpus. In our local BLAST search of OpenGenome2, we found more than 500 sequences similar to ΦX174 (**Supplementary Table 2**), compared with only nine similar to MS2. This difference is partly expected because Microviridae (ssDNA bacteriophages) are more abundant in current sequence repositories than phages with small RNA genomes, largely as a result of methodological ease.

At the genome-wide level, Evo2 again displayed broad sequence-level regularities. As in MS2, nonsynonymous changes received lower SLL scores than synonymous ones and stop codons lower still **(Fig. 4**). Reverse-complement scoring reduced the elevated variability at the 5′ end expected from limited upstream context in autoregressive scoring, while leaving the broader scoring pattern largely unchanged (**Fig. S5**). Interestingly, stop codons in genes *E* and *K*, both of which are embedded within other genes, were not heavily penalized, similar to the observation in the MS2 lysis region **(Fig. 4, Fig. S6-S8**). This suggests that overlapping genes may represent a particular challenge to Evo2.

**Figure 4.**
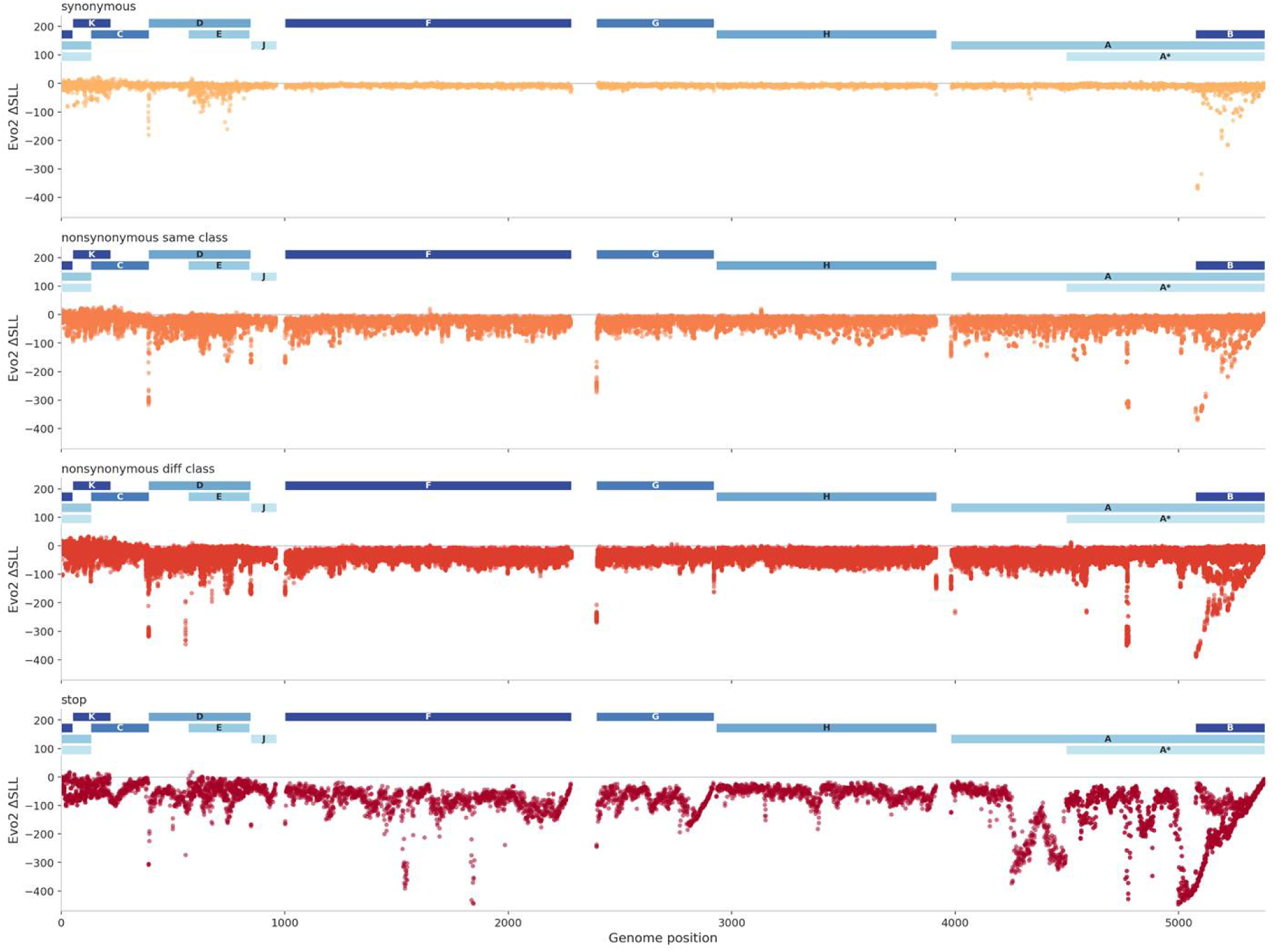
Genome-wide Evo2 scoring of codon substitutions in ΦX174. For each annotated coding position in the ΦX174 reference genome, every codon was substituted with each of the 63 alternative codons and scored with Evo2 7B. The y-axis shows ΔSLL, defined as the sequence log-likelihood of the mutant genome minus that of the reference genome; more negative values indicate variants that Evo2 considers less plausible than the reference. The x-axis shows genome position in plus-strand coordinates. Variants are separated into four effect classes: synonymous substitutions, nonsynonymous substitutions that preserve amino-acid class, nonsynonymous substitutions that cross amino-acid class, and nonsense substitutions introducing stop codons. Amino-acid classes were defined as nonpolar, polar, basic, and acidic, with tyrosine and glycine each treated as their own class. In contrast to MS2, the genome-wide ΔSLL landscape in ΦX174 did not show a comparably localized scoring irregularity.

The relationship between ΔSLL and *E. coli* codon usage was approximately three times stronger than in MS2 (ρ=0.240, p=1.7×10^−77^, n=5840; **Table 4**), and the correlation was significant across ten of the eleven annotated genes. Even so, codon usage explained less than 6% of the rank variance in synonymous codon scores, indicating that most of the Evo2’s codon-level preferences were driven by features other than simple host usage. ΔSLL did not correlate with codon usage in gene *K* (ρ=0.015, p=0.85), perhaps because *K* overlaps with other genes, so a synonymous change in *K* is often nonsynonymous in another gene. Thus, as in MS2, Evo2 distinguished among synonymous codons in ways that were only partly attributable to simple host codon preference.

**Table 4.** Evo2’s synonymous-codon preferences in ΦX174 are more aligned with *E. coli* codon usage than in MS2, but host usage still explains only a limited fraction of the variation. For each gene, the table reports the number of synonymous codon variants evaluated (“count”), together with the Spearman correlation between ΔSLL and the relative frequency with which each codon is used by *E. coli* within its synonymous set. Because ΦX174 depends on *E. coli* translation machinery, host codon usage provides one biologically plausible source of selective pressure on synonymous sites. Positive correlations therefore indicate that codons more frequently used by *E. coli* tend to receive higher Evo2 plausibility scores.

| GENE | COUNT | SPEARMAN<br>CORR. | P-VAL |
| --- | --- | --- | --- |
| A | 1254 | 0.235 | 3.38E-17 |
| A* | 864 | 0.214 | 2.16E-10 |
| B | 266 | 0.321 | 8.72E-08 |
| C | 237 | 0.165 | 0.0107 |
| D | 385 | 0.354 | 8.48E-13 |
| E | 269 | 0.153 | 0.0117 |
| F | 1060 | 0.292 | 2.49E-22 |
| G | 452 | 0.235 | 4.11E-07 |
| H | 783 | 0.31 | 6.01E-19 |
| J | 106 | 0.526 | 6.78E-09 |
| K | 164 | 0.015 | 0.853 |
| ALL<br>GENES | 5840 | 0.24 | 1.72E-77 |

### ΔSLL predicts viable from nonviable ΦX174 codons, but much of the signal is driven by nonsense mutations

We next determined whether Evo2 performed better in a DNA phage benchmark than it did in MS2. For this comparison, we used the recent ΦX174 gene *G* dataset of Van Leuven et al. [7], in which mutations were assessed in the context of reconstructed whole phage genomes rather than isolated gene expression. This provides a more uniformly generated benchmark for viability and complements the MS2 lysis dataset, where function was measured in a gene-isolation context. Specifically, the Van Leuven study built a large mutant set for gene *G*, the 172-aa spike/capsid protein, and experimentally measured whether each mutation yielded viable phage. Because their analysis was reported at the amino acid level rather than the codon level, some codons remained ambiguous in our reconstruction of the dataset. For the analyses here, we classified a codon as viable if it was recovered in the plaque pool or from site-directed mutagenesis or was the wild-type codon, and as nonviable if it introduced a stop codon, produced no plaques when constructed, or encoded an amino acid the authors reported as nonviable. This yielded 611 viable codons (590 viable mutant codons plus 21 wild type codons), 634 nonviable codons, and 99 codons of undetermined viability. Notably, unlike the MS2 lysis-gene benchmark, the ΦX174 gene *G* benchmark does not reside in an unusually dense overlapping regulatory context and did not exhibit stop codon scoring irregularities (**Fig. S6H, S7**).

ΔSLL showed a more favorable pattern for ΦX174 compared to MS2. Viable codons tended, on average, to receive higher scores than nonviable codons (p=9.1×10^−23^, AUROC = 0.66, n=1224; **Fig. 5A**). However, the distributions overlapped substantially. When we separated nonviable codons by mutation type, it became clear that most of the discrimination was driven by nonsense variants rather than by more subtle missense effects. Stop-codon mutations were strongly separated from viable codons (AUROC = 0.95, p=4.2×10^−32^, n=63), whereas the missense-only comparison was much weaker (AUROC = 0.63, p=1.5×10^−14^, n=571). Codons of undetermined viability did not differ from viable codons (AUROC = 0.55, p=0.11, n=99).

**Figure 5.**
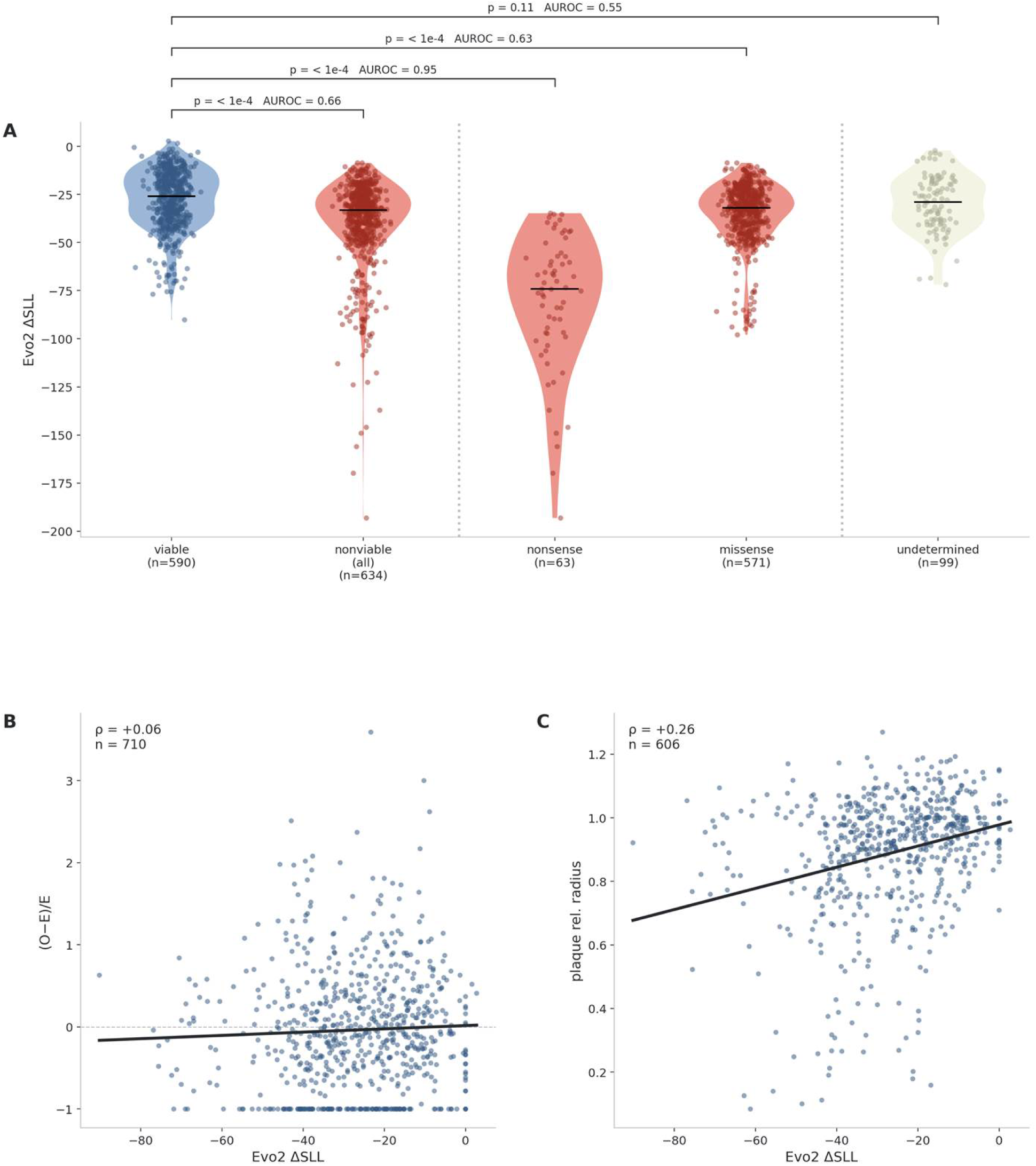
Evo2 ΔSLL shows limited phenotype signal in the ΦX174 gene *G* benchmark. **(A)** ΔSLL distributions for codons classified as viable, nonviable, or of undetermined viability in the Van Leuven et al. (2025) gene *G* dataset. The nonviable class is additionally partitioned into nonsense and missense variants to show that much of the separation from viable codons is driven by premature stop mutations. Statistical comparisons are benchmarked against the viable class. **(B)** Relationship between ΔSLL and the enrichment-like metric (observed−expected)/expected) derived from the starting library and recovered plaque pools in Van Leuven et al. (2025). Each point represents a codon variant; the line shows the fitted trend. **(C)** Relationship between ΔSLL and plaque radius for recovered gene *G* variants from Van Leuven et al. (2025). Each point represents a codon variant; the line shows the fitted trend.

We also assessed whether ΔSLL tracked two more quantitative outcomes reported in the Van Leuven study: enrichment of each variant relative to its starting library frequency and plaque size [7]. ΔSLL did not correlate with enrichment, measured as (observed−expected)/expected) (ρ=0.059, p=0.12, n=710; **Fig. 5B**). It did show a modest positive correlation with plaque size (ρ=0.263, p=4.7×10^−11^, n=606; **Fig. 5C**), suggesting that Evo2 captures some information relevant to stronger or weaker viable phenotypes even when it does not cleanly separate all viable from nonviable variants.

ΔSLL was also strongly associated with how many nucleotides separated a mutant codon from the reference codon (ρ=−0.349, p=3.5×10^−39^, n=1323; **Fig. S9A**). Codons one nucleotide away from the reference generally scored higher than codons two or three nucleotides away. This effect was stronger than either of the direct phenotype correlations and depends only on how different the sequence looks from the reference, not on the biological consequence of the substitution. To control for this, we repeated the enrichment and plaque-size correlations within groups of codons differing by one, two, or three nucleotides from the reference (**Fig. S9B,C**).

Within these strata, ΔSLL retained weak but detectable correlations with enrichment within the one-(ρ = 0.193, p = 0.0247, n = 136) or two-(ρ = 0.143, p = 0.0111, n = 317) nucleotide difference groups, but not in the group where three nucleotides differ from the reference (ρ = 0.0361, p = 0.581, n = 236; **Fig S9B**). ΔSLL’s correlation with plaque size persisted at two (ρ = 0.256, p = 1.7 × 10□□, n = 276) or three (ρ = 0.237, p = 7.8 × 10□□, n = 198) substitutions, but not one (ρ = 0.144, p = 0.12, n = 115; **Fig S9C**). Taken together, these analyses suggest that Evo2’s signal in ΦX174 is real but limited: it captures some biologically meaningful structure, but that structure remains entangled with simple reference-distance effects.

### Models of ΦX174 intermediate embeddings

Because raw ΔSLL showed at least modest phenotype signal in ΦX174, we next asked whether Evo2’s intermediate embeddings carried additional information beyond sequence likelihood alone. As in MS2, we evaluated embeddings from all 32 model blocks under multiple pooling schemes and trained simple supervised models to predict viability. The best-performing representation came from block 3 with site pooling, although the variation across configurations was noisy enough that little should be read into the specific winning layer (**Fig. 6A**). Under this best configuration, all models outperformed ΔSLL (**Fig. 6B,C**, **Table 5**). UMAP visualization again failed to show clear phenotype-separated clusters in the first two dimensions (**Fig. 6D**).

**Figure 6.**
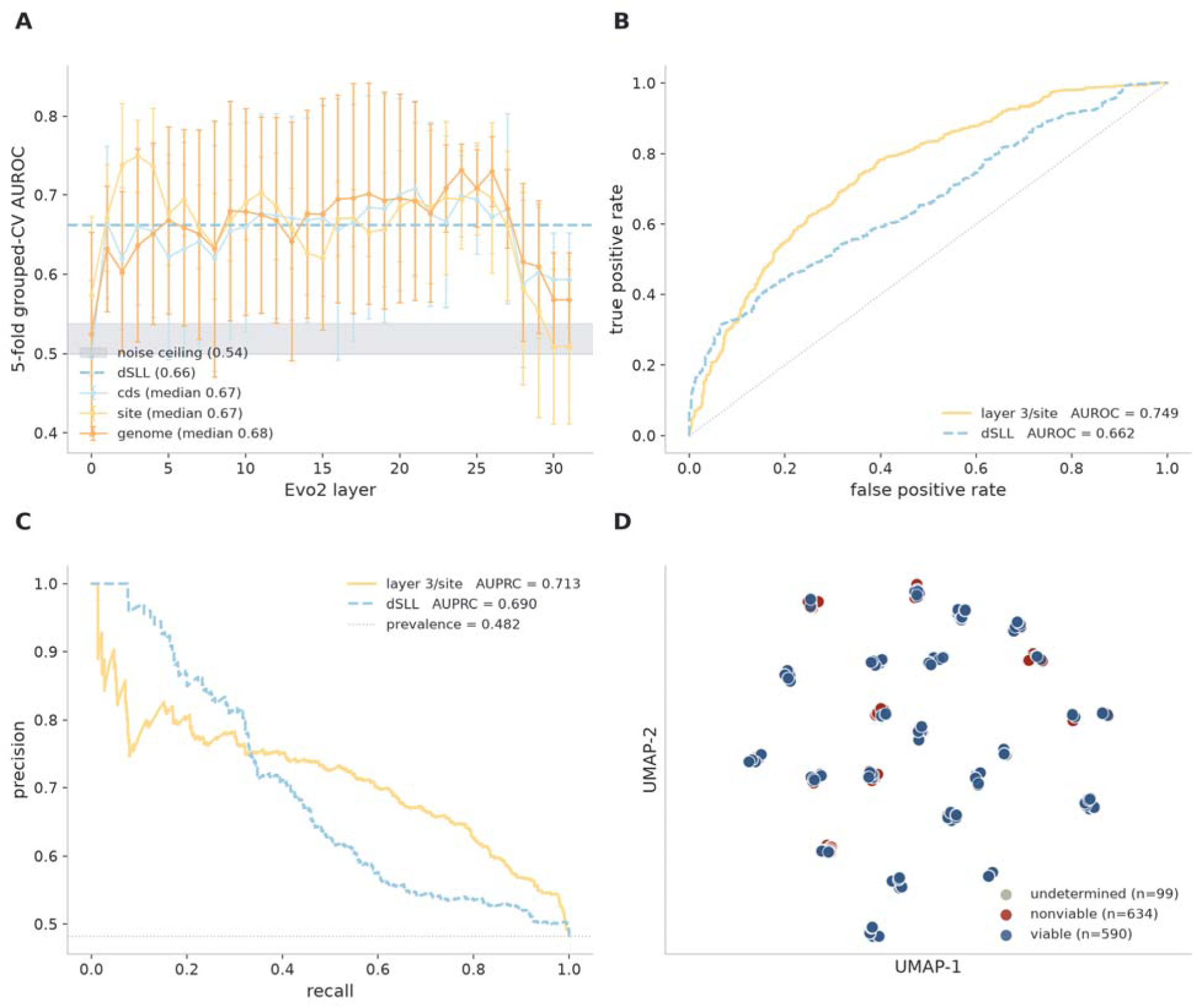
Intermediate Evo2 embeddings provide limited and unresolved phenotype signal in the ΦX174 benchmark. **(A)** AUROC from grouped five-fold cross-validation across Evo2 layers for the three pooling schemes tested: coding sequence (CDS), mutated site, and whole genome. Grouping was performed by mutation site so that different substitutions at the same position were always kept in the same held-out subset. **(B)** Receiver operating characteristic (ROC) curve for the best-performing model configuration, block 3 with site pooling (yellow), compared with ΔSLL used directly as a score (blue dashed). **(C)** Precision-recall curve for the same comparison as in panel B. The gray dotted line indicates class prevalence. **(D)** UMAP projection of the best-performing embedding representation (block 3, site pooling). Each point represents a codon variant; blue points indicate viable variants and red points indicate nonviable variants. As in MS2, the projection does not cleanly separate phenotype classes.

**Table 5.**
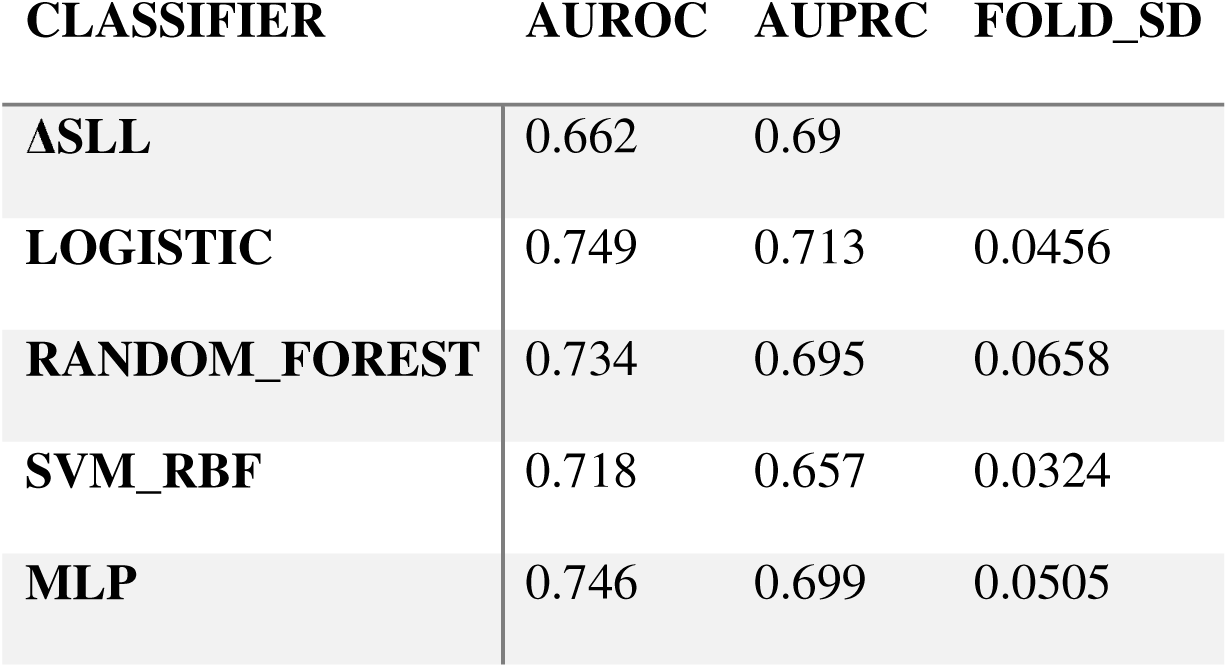
Classifier comparison on Φ*X*174 embeddings (block 3, site pooling). Grouped five-fold cross-validation; 590 viable and 634 nonviable variants across 21 sites. Columns as in Table 2.

| CLASSIFIER | AUROC | AUPRC | FOLD_SD |
| --- | --- | --- | --- |
| ΔSLL | 0.662 | 0.69 |  |
| LOGISTIC | 0.749 | 0.713 | 0.0456 |
| RANDOM_FOREST | 0.734 | 0.695 | 0.0658 |
| SVM_RBF | 0.718 | 0.657 | 0.0324 |
| MLP | 0.746 | 0.699 | 0.0505 |

We then asked whether this apparent embedding advantage reflected genuine phenotype information or simpler properties of the dataset. To do so, we repeated the covariate analysis used for MS2, here adding the number of nucleotide differences from the reference codon as an explicit variable alongside genomic position and stop status (**Table 6**). Covariates alone were not as strong predictors of phenotype as in the MS2 case (AUROC = 0.629 for logistic regression, AUROC = 0.671 for random forest). Adding embeddings improved the performance of both classifiers (AUROC = 0.757 for logistic regression, AUROC = 0.724 for random forest), suggesting that they do provide predictive signal beyond the simple covariates, but still do not cleanly separate viable and nonviable mutants.

**Table 6.** On Φ*X*174, covariates and embeddings provide partially overlapping but not decisive phenotype signal. Columns as in Table 3; covariates here are genomic position, stop status, and the number of nucleotide substitutions separating the codon from the reference, and embeddings are block 3 with site pooling. AUROC is oriented so that a value above 0.5 means viable codons score higher.

| CLASSIFIER | FEATURES | N_FEATURES | AUROC |
| --- | --- | --- | --- |
| LOGISTIC | site/position only | 1 | 0.429 |
|  | is_stop only | 1 | 0.477 |
|  | nt_changes only | 1 | 0.582 |
|  | all covariates | 3 | 0.629 |
|  | embeddings only | 4096 | 0.749 |
|  | all covariates + emb | 4099 | 0.757 |
| RANDOM FOREST | site/position only | 1 | 0.599 |
|  | is_stop only | 1 | 0.477 |
|  | nt_changes only | 1 | 0.582 |
|  | all covariates | 3 | 0.671 |
|  | embeddings only | 4096 | 0.734 |
|  | all covariates + emb | 4099 | 0.724 |

We therefore treat the ΦX174 model results as suggestive, but not decisive, with respect to positional and reference-distance confounding. Compared with MS2, ΦX174 offers a more favorable and less anomalous benchmark, but even here the signal falls short of a simple interpretation in which model scores or embeddings can be taken as direct readouts of biological function.

## DISCUSSION

### Why MS2 and ΦX174 behave differently

A central lesson from this study is that current genome language models can recover expected mutation-class (e.g., synonymous vs stop) and codon-specific changes (polar vs non-polar) in compact phage genomes without yet delivering reliable functional mutational-effect prediction. Across both phages, synonymous mutations tended to remain closer to the reference in score, whereas nonsynonymous mutations produced a broader range of model responses, and nonsense mutations were often strongly penalized **(Fig. 1**, **Fig. 4, Fig. S3, Fig. S7)**. Evo2 produced varying ΔSLL scores for synonymous mutations occurring at the same site, with only a limited fraction of that variation explained by host codon usage **(Table 1**, **Table 4)**, suggesting that the model has learned sequence constraints beyond simple codon-bias effects. However, the failure to strongly penalize stop codons in the MS2 lysis gene suggests Evo2 may struggle to process overlapping coding regions. Specifically, Evo2 treated many truncating lysis mutations as near-reference sequences even though they should be among the clearest loss-of-function changes in the benchmark **(Fig. 1, Fig. S2C, Fig. S3D, Fig. S4)**.

MS2 and ΦX174 are both small, tractable, and historically rich systems, but they challenge genome language models in different ways. MS2 imposes RNA-level constraints beyond protein coding, including regulatory RNA structures that control translation and replication. Accumulation of coat protein represses replicase translation through binding to a defined RNA operator, and the timing and stoichiometry of gene expression depend on folding transitions embedded in the genome itself. In this setting, even sequence changes that appear silent at the protein level can plausibly disrupt RNA structure, translation control, or genome packaging. This makes MS2 a particularly demanding test of whether a model is learning biologically meaningful constraints or merely sequence plausibility.

ΦX174 was used to provide a less challenging DNA-phage benchmark. Its genome is DNA rather than RNA, the gene under direct mutational study does not lie within the same densely overlapping regulatory context as MS2 lysis, and the benchmark mutations were assessed in reconstructed whole phage genomes rather than isolated gene expression. These differences likely contribute to why MS2 lysis behaved as a strong stress-test case, whereas ΦX174 provided a more favorable, though still limited, benchmark. Taken together, the two phages offer complementary tests of model behavior across distinct layers of biology.

### Phage function as a benchmark of model performance

This work is not simply the evaluation of a single model checkpoint, software version, or transient model state, but the establishment of what we view as a reusable benchmarking framework. Specifically, leveraging well defined phage genomes, and associated experimental datasets to test, validate, and improve GLMs. A major strength of this approach is the use of published experimental mutation datasets that are not typically represented as structured genotype-to-phenotype records in sequence repositories. These data are valuable precisely because they were costly to generate, are experimentally grounded, and encode viable versus nonviable outcomes that can be used immediately for model evaluation without first generating a large new wet-lab training dataset. They also prevent the benchmark from collapsing onto whatever is easiest to extract from modern sequence databases, which may not reflect the full depth or difficulty of experimentally validated genotype-to-phenotype knowledge.

This study is motivated by a broader long-term vision, which we refer to as “prompt-to-phage” design [9]: a researcher specifies a biological objective, and an AI system proposes novel, chimeric, or edited phage genomes that can be tested experimentally. A recent study showed that AI workflows can generate viable ΦX174-like phages [5]; however, it remains unclear how much of that success reflects model-derived biological understanding versus strong human guidance or whether similar approaches could succeed for other phages. Our work addresses a different question, but one that would likely need to be resolved before a broadly useful prompt-to-phage engine could be realized: whether current genome language models can reliably predict the effects of simple sequence changes in compact phage genomes. In that sense, this study introduces an early proficiency checkpoint on the path toward prompt-to-phage design that can be benchmarked systematically.

Raw sequence likelihood alone was insufficient in MS2 lysis benchmark and only modestly informative in ΦX174. Embedding-based models improved apparent prediction relative to likelihood scoring, but those improvements required careful interpretation because simple covariates could explain or complicate the signal **(Fig. 2**, **Fig. 3**, **Table 2**, **Table 3**, **Table 5**, **Table 6)**. Thus, intermediate model representations may be useful for future predictive workflows, but they should not be treated as direct evidence of biological ‘understanding’ unless they outperform appropriate covariate baselines. Overall, we believe this evidence suggests that more complex agentic workflows may be required to advance a true ‘prompt-to-phage” system, in which genome language models may serve as a tool for an agent, but functional data extracted from the literature will be required for robust functionally aware ‘designer’ phages.

### Limitations

A key limitation of this study is that the MS2 mutation dataset used here cannot be treated as perfect genome-wide ground truth. Mutations that abolish lysis in the Chamakura dataset [8] are likely to remain strongly deleterious in a whole-genome context. In contrast, permissive mutations identified when the lysis gene is assayed in isolation cannot be assumed to remain permissive once placed back into the intact genome, where the same region also encodes overlapping reading frames and RNA regulatory features. However, it does not invalidate the use of MS2 as a benchmark. First, the MS2 result does not depend only on how one interprets the permissive class. Even mutations that should clearly block function in this assay context (e.g., lysis-gene stop codons) were not consistently scored as strongly deleterious by Evo2 **(Fig. 2A&C; Fig. S2C; Fig. S3D; Fig. S4)**. Second, the value of the MS2 dataset here is not that it provides perfect whole-genome truth, but that it exposes a difficult, biologically dense benchmark. Third, we paired MS2 with a cleaner and more systematic ΦX174 benchmark precisely to reduce the risk that the study would hinge on a single difficult dataset. Read this way, the MS2 analysis is useful not as a final adjudication of genotype-to-phenotype prediction, but as a stringent benchmark for future studies and a source of testable hypotheses. A natural next step is to use our previously reported cDNA platform, to empirically test, at genome scale, the boundary conditions highlighted here predicted to be most-tolerated and least-tolerated. This approach would directly expand the genotype-to-phenotype corpus needed to evaluate the current findings and to refine future models.

As discussed above, the ΦX174 dataset is easier to interpret and more closely aligned with the model’s training substrate: it is a DNA phage, and is more richly represented in the training corpus. Furthermore, mutations in gene *G* were assessed in the context of reconstructed whole phage genomes, and *G* is not embedded within the overlapping gene architecture seen in MS2. Even with these advantages, Evo2 did not yield a high-fidelity predictor of phage fitness. In our results, ΦX174 showed that Evo2 can recognize the clearest failure cases, particularly mutations that introduce premature stop codons **(Fig. 5A; Fig. S6–S8)**. Beyond those obvious disruptions, however, it did not provide a consistently reliable predictor of whether a mutant phage would remain viable **(Fig. 5A–C; Fig. S9)**. Future work should therefore aim to generate a large volume of genome-wide MS2 mutation-fate data, ideally at single-nucleotide resolution, while using ΦX174-like datasets as complementary benchmarks.

An addition limitation of the present study is that we evaluated Evo2 7B rather than the largest available variants. We therefore do not interpret these results as defining the ceiling of genome language model performance on phage mutation prediction. Larger models may behave differently and may improve the results reported here. At the same time, our choice was deliberate. Evo2 7B represents a level of computational access that is more realistic for many research groups, where GPU availability, queue time, and software stability can constrain routine use of frontier-scale models. The present results are therefore informative not only about model behavior, but about what can be achieved with tools that are realistically accessible for iterative biological benchmarking.

## CONCLUSION

Phages are useful proving grounds for testing the readiness and credibility of genome language models in biological prediction because they are compact, experimentally tractable, and provide a rapid bridge between sequence and function. Well-characterized model phages also offer a comparatively safe system for iterative prediction-testing benchmarks while still exposing real biological constraints. At the same time, phages are not trivial systems. Like more complex forms of life, they expose constraints on genome organization, molecular function, gene regulation, and replication. They therefore provide a meaningful test of whether models are learning biological relationships or primarily sequence plausibility. The framework established here provides a durable way to assess future model readiness. As genotype-to-phenotype data accumulate and models gain richer biological context, phages offer a practical way to identify when prediction has become reliable enough to warrant trust, and when it has not.

## STAR METHODS

### Data curation

#### Selection of datasets

We used SPARKIT (https://sparkit.science) to search the literature for bacteriophage mutational analyses suitable for this study. We prioritized the datasets of Chamakura et al. [8] and Van Leuven et al. [7] because of the small genomes of the phages studied, the availability of nucleotide-level sequence data linked to phenotypic measurements, and the large number of variants tested.

#### Reference genomes

Recent work from our laboratory showed that the MS2 reference sequence (NC_001417.2) is nonviable and identified the minimal corrections required to restore viability. For the MS2 benchmark analyses, we used the corrected reference background without the additional maturase D241H substitution that arose during the cDNA reconstruction study. This choice kept the benchmark sequence as close as possible to the original reference genome and avoided conflating baseline reference correction with the behavior of a derived incidental SNP. Neither the reference sequence nor the nearest functional isolate tested here carried this change. Although the D241H substitution proved compatible with phage production in our system, and comparison to other genomes suggests that this position is not absolutely constrained to aspartate, the overwhelming majority of related isolates retain D241. For benchmarking purposes, we therefore treated the D241H-free corrected sequence as the leanest and most interpretable rescued reference background [9]. Chamakura et al. (2017) expressed only the MS2 lysis gene and did not publish the nucleotide sequence of the reference they used. However, the amino acid sequence of the lysis gene matched both the original reference genome (NC_001417.2) and the corrected reference genome ([insert accession]), so we used the corrected functional sequence ([insert citation]) as the MS2 reference background for all analyses.

Van Leuven et al. [7] stated that they used the ΦX174 strain with accession AF176034.1, and we used the same reference in our analysis. At the time of analysis, the Van Leuven et al. dataset was available as a public preprint under revision; correspondence with the corresponding author indicated that the raw data and viable/nonviable mutation calls were not expected to change materially in the revised version.

We annotated the corrected MS2 sequence using the NC_001417.2 RefSeq GFF3 file and the ΦX174 AF176034.1 sequence using the NC_001422.1 RefSeq GFF3 file. In both cases, the experimental reference aligned one-to-one with the corresponding RefSeq record without rotation, allowing coordinates to transfer directly.

### OpenGenome2 homology search and hit sequence retrieval

OpenGenome2 [1] nucleotide FASTA files were downloaded from the Hugging Face repository arcinstitute/opengenome2; the successfully completed download finished on June 12, 2026. A local nucleotide BLAST database was constructed using makeblastdb from BLAST+ v2.17.0 with nucleotide database type (-dbtype nucl) and sequence-identifier parsing enabled (-parse_seqids). Before database construction, pipe characters in sequence identifiers were replaced with underscores, identifiers longer than 50 characters were truncated, and numeric suffixes were added when necessary to maintain uniqueness.

The reference genomes described above were searched against the local OpenGenome2 database using BLASTN v2.17.0 with an E-value threshold of 1×10^−5^, a maximum of 500 reported target sequences per query, and 16 threads. Results were written in XML format for downstream processing.

The workflow was managed using Snakemake v9.23.1, and BLAST XML output was parsed using Biopython v1.87. Detailed hit tables contained one row per high-scoring segment pair (HSP) and reported the query identifier, normalized database accession, hit description, E-value, bit score, percent identity, and query coverage. Percent identity was calculated as the number of identical aligned positions divided by the HSP alignment length. Query coverage was calculated from the aligned query span relative to the full query length.

A separate summary table was generated for each unique database hit across all of its HSPs. Overlapping or adjacent query intervals were merged to calculate combined query coverage, and the summary reported the number of HSPs, covered query intervals, best E-value, best bit score, and HSP-level percent-identity statistics. Duplicate accessions were removed, and the full nucleotide sequence for every unique reported hit was retrieved from the local database using blastdbcmd and written to FASTA format. Because BLAST output was limited to 500 target sequences per query, the retrieved FASTA files contain all unique hits reported within that limit.

### Variant enumeration

Two enumerations were generated for each genome. The single-nucleotide path substituted every position with each of the three alternative bases. The codon path substituted every codon within each annotated CDS with each of the 63 alternative codons. Codon substitutions in overlapping reading frames are enumerated once per gene, so the same mutated genome can appear under more than one gene annotation. For simplicity, only the codon-level results are shown in the main text, as these contain all of the information in the single-nucleotide scans and more. The single-nucleotide enumerations were nonetheless generated and used in the analysis pipeline, including the scripts that produce **Fig. 2**.

### Evo2 scoring

Each enumerated variant was scored with Evo2 7B, checkpoint arcinstitute/evo2_7b, obtained from the Hugging Face Hub through the evo2 package (v0.5.5). Its internal architecture (StripedHyena 2, implemented in the vtx package, v1.0.8) was not modified, and no parameter of the model was trained or updated at any point. Evo2 is an autoregressive model: starting from the 5′ end, it assigns a probability to each nucleotide based on the sequence that precedes it. The sequence log-likelihood (SLL) is the sum of the logarithms of those probabilities. For every variant, the complete mutated genome was scored rather than a local window around the substitution. The unmutated reference genome was scored once under identical settings, and the reported quantity is ΔSLL = SLL(mutant genome) − SLL(reference genome), reported in nats. Negative values indicate that the model considers the mutant less likely than the reference. Because both sequences are scored end-to-end and differ at one to three positions, ΔSLL combines the direct change in log-likelihood at the substituted position with the indirect change at every downstream position, which is conditioned on altered upstream context. All analyses below use rankings or differences of ΔSLL, so neither the unit nor the choice to sum rather than average across positions affects the conclusions.

Each genome was scored under two settings. The first used a single forward pass over the plus strand. The second averaged the forward pass with a pass over the reverse complement of the same sequence, using Evo2’s average_reverse_complement option. A forward pass conditions each position only on the sequence upstream of it, so positions near the 5′ end are scored with limited context, whereas a reverse-complement pass supplies context to those positions but reads a positive-sense RNA genome in the opposite direction from translation and replication.

Identical mutated sequences were scored once and the result joined back to every variant row that produced them. Two sequences were treated as identical when their SHA-256 digests matched. This matters only for the codon enumeration, where overlapping reading frames can cause the same mutated genome to be enumerated under two genes. Scoring used a single NVIDIA H100 with 80 GB of memory, with batch sizes of 32 for MS2 and 16 for ΦX174.

### Variant annotation

Scored variants were annotated against the GFF3 files by locating each position within a CDS, determining the affected codon and its position within the codon, and translating the reference and alternative codons. Coding changes were assigned to four mutually exclusive effect types: synonymous, nonsynonymous within an amino-acid class, nonsynonymous across amino-acid classes, and nonsense. Amino-acid classes were defined as nonpolar (A, V, I, L, M, F, W, P), polar (S, T, C, N, Q), basic (K, R, H), and acidic (D, E). Tyrosine and glycine were each treated as their own class rather than grouped with the others.

### Published phenotype data

#### MS2 lysis protein L

Variant-level lysis phenotypes were taken from supplementary table S2 of Chamakura et al. (2017). That table lists sequenced isolates rather than distinct variants: 139 rows collapse to 83 distinct single-nucleotide substitutions, with one substitution recovered up to 12 times. Isolates were deduplicated on position, reference base, and alternative base; no substitution carried conflicting lysis calls across its isolates. Each variant therefore carries a binary lysis phenotype and a western blot call for L protein (present, absent, or not determined). Variants were additionally stratified by reading frame, because the lysis gene overlaps the coat and replicase genes and only a minority of substitutions fall in the lysis gene alone.

#### ΦX174 gene G

Codon-level viability data were taken from Van Leuven et al. [7], rebuilt from the per-plaque Sanger calls in the repository rather than from the summary sheets in the supplemental data. A codon was called viable if it was the wild-type codon, was recovered at least once in the plaque pool, or was constructed by site-directed mutagenesis and produced plaques. A codon was called nonviable if it introduced a stop codon, if it was constructed and produced no plaques, or if it encoded an amino acid at a position the source publication called nonviable. Codons meeting neither definition were treated as undetermined and were not used to train the model.

### Embedding extraction and linear models

As Evo2 reads each base, it builds an internal representation, or embedding, of that position: a 4096-dimensional vector per nucleotide. A full-genome pass therefore yields a 3569×4096 matrix for MS2 and a 5386×4096 matrix for ΦX174. Each of the model’s 32 successive processing blocks produces its own such representation, with the final probabilities read out from the last block through a linear layer. We extracted embeddings from each block and trained logistic regression models using the experimental phenotype as the target variable: lysis function for MS2 and viability for ΦX174.

For each variant, we extracted representations from all 32 blocks under two sequence contexts: a local pass spanning the affected coding sequence (gene *L* for MS2 or gene *G* for ΦX174) plus 500 bases of flanking sequence on each side, and a whole-genome pass covering the entire mutated genome. Because the model requires a single vector per variant rather than one vector per nucleotide, the per-position embeddings were averaged before classification. We compared three pooling schemes. In the CDS scheme, embeddings were averaged across the full coding sequence from the local pass. In the site scheme, embeddings were averaged only across the three positions of the mutated codon from that same local pass. In the genome scheme, embeddings were averaged across the entire genome from the whole-genome pass.

CDS and site therefore use the same underlying local representations and differ only in the width of the window averaged. The three pooling schemes ask whether any phenotype signal is local to the substituted codon, distributed across the gene, or detectable in the model’s representation of the genome as a whole.

Each of the 96 combinations of layer and pooling (32 blocks × 3 pooling schemes) was evaluated with logistic regression using scikit-learn 1.7.2. Features were standardized to zero mean and unit variance. An L2 penalty (C = 1.0) and an increased iteration limit of 5000 were used; all other settings were left at default (LogisticRegression(max_iter=5000, C=1.0)). Performance was measured by five-fold stratified grouped cross-validation: the data were split into five parts, the model was trained on four and tested on the fifth, and this was repeated so that every variant was predicted once by a model that had not seen it. Folds were grouped by codon site so that variants at the same site never appeared in both training and test partitions. This grouping is necessary because both datasets contain multiple substitutions at the same position; without grouping, the classifier could learn site identity rather than the effect of the substitution. All estimators were seeded so that rerunning the analysis reproduced the same results.

Taking the maximum of 96 configurations tends to overstate performance. However, the aim was to give Evo2 the most favorable representation possible rather than evaluate one dependent on an arbitrary layer or pooling choice. The median performance across layers within each pooling scheme is therefore also reported, as it is not inflated by post hoc selection of a single best configuration. To estimate the size of this inflation directly, the full 96-configuration sweep was repeated 50 times on random features containing no information about the phenotype. The resulting performance distribution was treated as a “noise ceiling,” and an observed maximum had to exceed that ceiling to be meaningful.

### Comparison of classifiers

To give Evo2 the best possible chance at phenotype prediction, we next asked whether its intermediate embeddings contained phenotype-relevant information that might not be recoverable with a simple linear model. We therefore evaluated three additional classifiers alongside logistic regression: a random forest, a support vector machine with a radial basis function kernel, and a multilayer perceptron with one hidden layer. All classifiers were applied to the same fixed embedding representation, namely the best-performing layer and pooling configuration identified by the logistic-regression sweep, and used functionality (for MS2) or viability (for ΦX174) as the target variable. All classifiers were fit on standardized features under the same grouped cross-validation scheme and were compared directly against ΔSLL used as a score without additional training. The purpose of this comparison was not to optimize predictive performance at all costs, but to ask a narrower question: if useful phenotype information is present in the embeddings, is it linearly accessible, or does it emerge only when a more flexible classifier is allowed to fit nonlinear patterns?

We deliberately did not perform an extensive hyperparameter search. Aggressive tuning would introduce a second layer of model selection on the same held-out data and would make it more difficult to interpret whether performance differences reflected real signal or simply extra optimization effort. Instead, we used standard implementations with only minimal changes needed for stability and convergence. Logistic regression used an L2 penalty (C=1.0) and an iteration limit of 5000, as above (LogisticRegression(max_iter=5000, C=1.0)). The random forest used 300 trees rather than the default 100 to reduce variance in the ensemble (RandomForestClassifier(n_estimators=300, random_state=0)). The support vector classifier used a radial basis function kernel with probability calibration enabled (SVC(kernel=’rbf’, probability=True, random_state=0). The multilayer perceptron used 128 hidden units and an iteration limit of 1000 to ensure convergence (MLPClassifier(hidden_layer_sizes=(128,), max_iter=1000, random_state=0)). All classifiers were implemented using scikit-learn 1.7.2.

### Confounding by variant covariates

A classifier can appear to perform well for reasons that have little to do with biological understanding. For example, it may learn that mutations in one region of a gene are usually harmful and mutations in another region are often tolerated, without learning anything specific about why a given substitution changes function. Grouping variants by site in cross-validation prevents the most direct form of this problem—training on one mutation at a site and testing on another mutation at that same site—but it does not eliminate a broader positional effect. A model can still learn that nearby sites tend to behave similarly. This is particularly relevant here because embeddings are computed from local or genome-scale sequence windows and therefore can implicitly encode where in the genome a mutation occurs, not just what the mutation is.

To test whether the model signal reflected true phenotype information or simpler properties of the dataset, we compared the embedding-based classifiers against models trained on basic covariates alone. These covariates were simple non-embedding features of each mutation: genomic position, whether the substitution introduced a stop codon, and, for ΦX174, the number of nucleotide differences separating the mutant codon from the reference. We evaluated each covariate set alone, in combination, and then concatenated with the embeddings, using the same classifiers and the same grouped cross-validation scheme as in the main model analysis. We interpreted embeddings as contributing information beyond a given covariate only if the embedding-based model outperformed the covariate model alone and if adding the covariates to the embeddings did not further improve performance.

### Dimensionality reduction

Two- and three-dimensional projections were also generated using Uniform Manifold Approximation and Projection (UMAP), a dimensionality-reduction method that places high-dimensional data into a low-dimensional space for visualization. UMAP was applied to the layer and pooling configuration selected by the model sweep using standardized features, 15 neighbors, a minimum distance of 0.1, Euclidean distance, and a fixed random seed. These projections were used only for qualitative visualization of the embedding structure and were not interpreted quantitatively.

### Statistical analysis

Comparisons between two groups used the two-sided Mann-Whitney U test. Effect sizes are reported as AUROC. Intuitively, AUROC can be interpreted as the probability that a randomly chosen functional variant receives a higher score than a randomly chosen nonfunctional variant. A value of 0.5 indicates complete overlap between the two groups, such that the score carries no information about phenotype. Values above 0.5 indicate that functional variants tend to score higher, whereas values below 0.5 indicate that the model ranks the two groups in the opposite direction from expectation.

The AUROC is directly related to the Mann-Whitney U statistic. Specifically, it is the proportion of all functional-versus-nonfunctional pairs in which the functional variant scores higher, with ties counted as half a pair. This means that the p-value and the AUROC reported for a comparison derive from the same underlying rank statistic: the p-value tests whether the observed pair ordering is unlikely under the null hypothesis of no difference, while the AUROC reports the magnitude and direction of the separation itself. Because AUROC was also used to quantify classifier performance, group comparisons, model evaluations, and noise-ceiling analyses are all reported on a common and interpretable scale.

Comparisons across more than two mutually exclusive groups used the Kruskal-Wallis test, followed where significant by pairwise Mann-Whitney tests with Holm-Bonferroni correction across the family. Each pairwise comparison was re-ranked only within the two groups involved, preserving the same AUROC interpretation as in the two-group case. Nonparametric tests were used throughout because ΔSLL distributions were heavy-tailed and strongly influenced by nonsense variants. Correlations between ΔSLL and continuous phenotype measurements were assessed with Spearman rank correlation.

## Data and software availability

The analysis is composed of two Snakemake workflows (v9.23.1), one covering the OpenGenome2 sequence search and the other covering enumeration, scoring, annotation, phenotype joins, embedding extraction, probing, and figure generation as a single dependency graph, so that every reported figure and table is reproducible from the reference genomes and the published phenotype tables. Jobs were dispatched to SLURM through snakemake-executor-plugin-slurm (v2.7.1); scoring and embedding extraction were the only GPU-requiring steps.

Analysis used Python 3.12 with Evo2 0.5.5, vtx 1.0.8, PyTorch 2.7.1 (CUDA 12.9), flash-attn 2.8.0.post2, scikit-learn 1.7.2, umap-learn 0.5.12, NumPy 2.4.6, pandas 2.3.3, SciPy 1.16.3, Biopython 1.87, and Matplotlib 3.10.8. A complete environment listing for each workflow is included in the repository as environment_versions.txt.

## CRediT AUTHOR CONTRIBUTIONS

Emily M. Layton: Conceptualization, data curation, formal analysis, funding acquisition, investigation, methodology, software, validation, visualization, writing—original draft, and writing—review and editing.

Michael L. Bernauer: Conceptualization, funding acquisition, investigation, methodology, and writing—review and editing.

Laura D. Weinstock: Conceptualization, formal analysis, methodology, supervision, validation, and writing—review and editing.

Eric M. Small: Conceptualization, data curation, investigation, methodology, project administration, resources, supervision, validation, visualization, and writing—review and editing.

Gina M. Geiselman: Visualization, writing—original draft, and writing—review and editing.

George Bachand: Conceptualization, validation, and writing—review and editing.

Jesse L. Cahill: Conceptualization, data curation, funding acquisition, investigation, methodology, project administration, resources, supervision, validation, writing—original draft, and writing—review and editing.

## ACKNOWLEDGEMENTS

We thank members of the Cahill lab for thoughtful discussion in support of this work. We thank Isabella Romano for a thoughtful technical review of this manuscript.

This work was supported by the Laboratory Directed Research and Development program at Sandia National Laboratories. Sandia National Laboratories is a multimission laboratory managed and operated by National Technology & Engineering Solutions of Sandia, LLC, a wholly owned subsidiary of Honeywell International Inc., for the U.S. Department of Energy’s National Nuclear Security Administration under contract DE-NA0003525.

E.M.L was supported by the Universities Research Association - Sandia Graduate Student Summer Fellowship and the Lawrence M. Blatt Biotechnology Internship Scholarship.

This paper describes objective technical results and analysis. Any subjective views or opinions that might be expressed in the paper do not necessarily represent the views of the U.S. Department of Energy or the United States Government

## DECLARATION OF INTERESTS

The authors declare no competing interests.

## Notes

### Competing Interest Statement

The authors have declared no competing interest.

