## Supplemental Data for "The phage stress test: a proving ground for genome language models in biological prediction"

**SUPPLEMENTAL FIGURES**

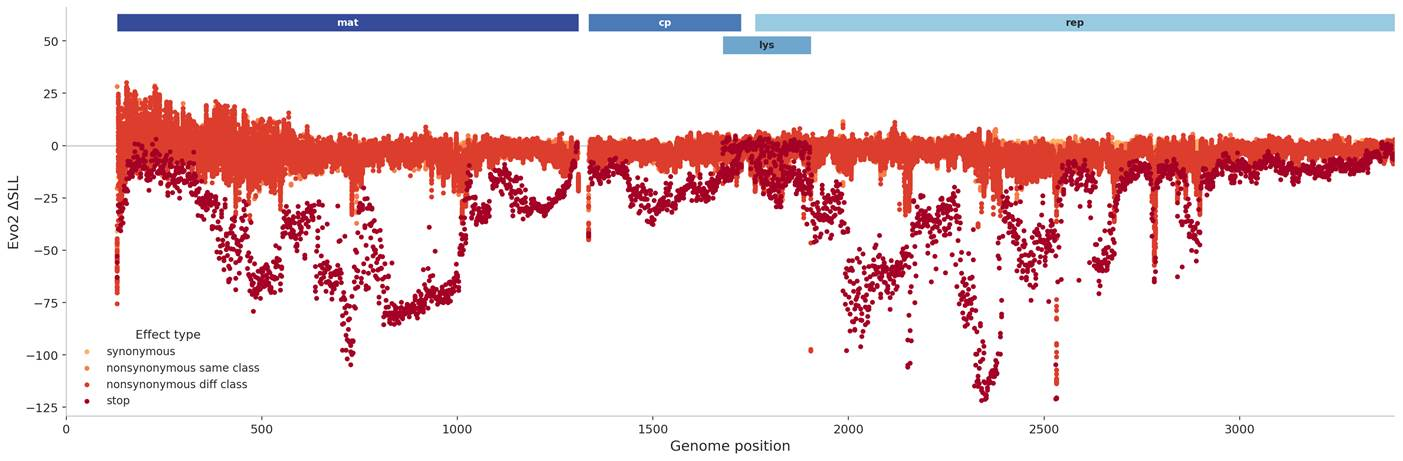

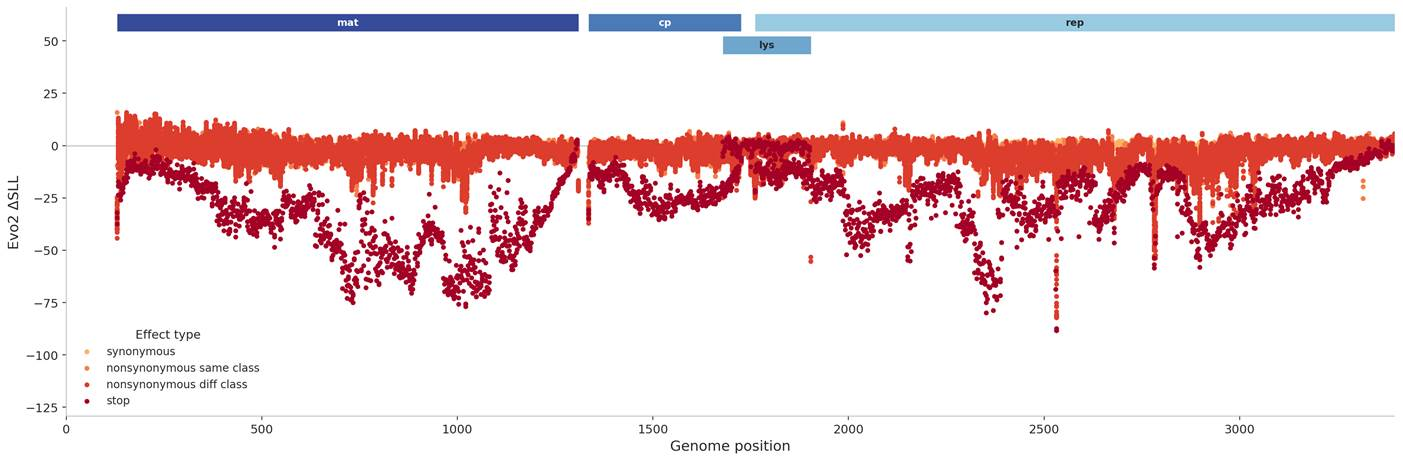

**Figure S1. Reverse-complement scoring reduces the 5′-end artifacts in MS2 genome-wide ΔSLL analysis.** Top, genome-wide MS2 codon-substitution scores calculated from the standard forward Evo2 pass. Bottom, the same analysis calculated using Evo2’s reverse-complement mode, in which the forward score and the reverse-complement score are averaged. Reverse-complement scoring reduces the elevated variability observed near the 5′ end of the genome in the forward-only analysis, consistent with this pattern arising from limited upstream context in autoregressive scoring rather than from a biological feature of the MS2 genome.

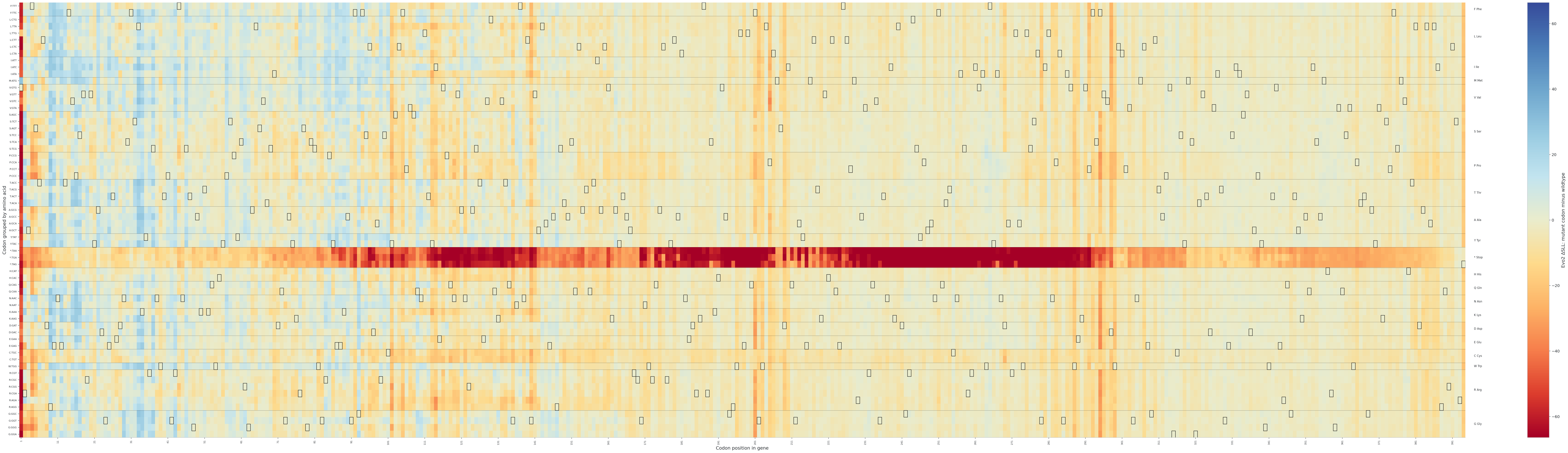

**Figure S2A. Codon-resolution heatmaps of Evo2** **ΔSLL scores across the MS2 maturase gene.** The y-axis shows the substituted codon, grouped by encoded amino acid and labeled by nucleotide triplet, and the x-axis shows codon position within the indicated gene. Each cell reports the ΔSLL of the full mutant genome after replacement of the indicated codon at the indicated site. Rows within each amino acid block are ordered by *E. coli* codon usage, with the most frequently used codon for that amino acid at the top and descending frequencies beneath. Boxes outline wildtype codons. The heatmap provides codon-level resolution of the genome-wide scoring patterns shown in Fig. 1 and allows localized inspection of tolerated and strongly disfavored substitutions within each gene.

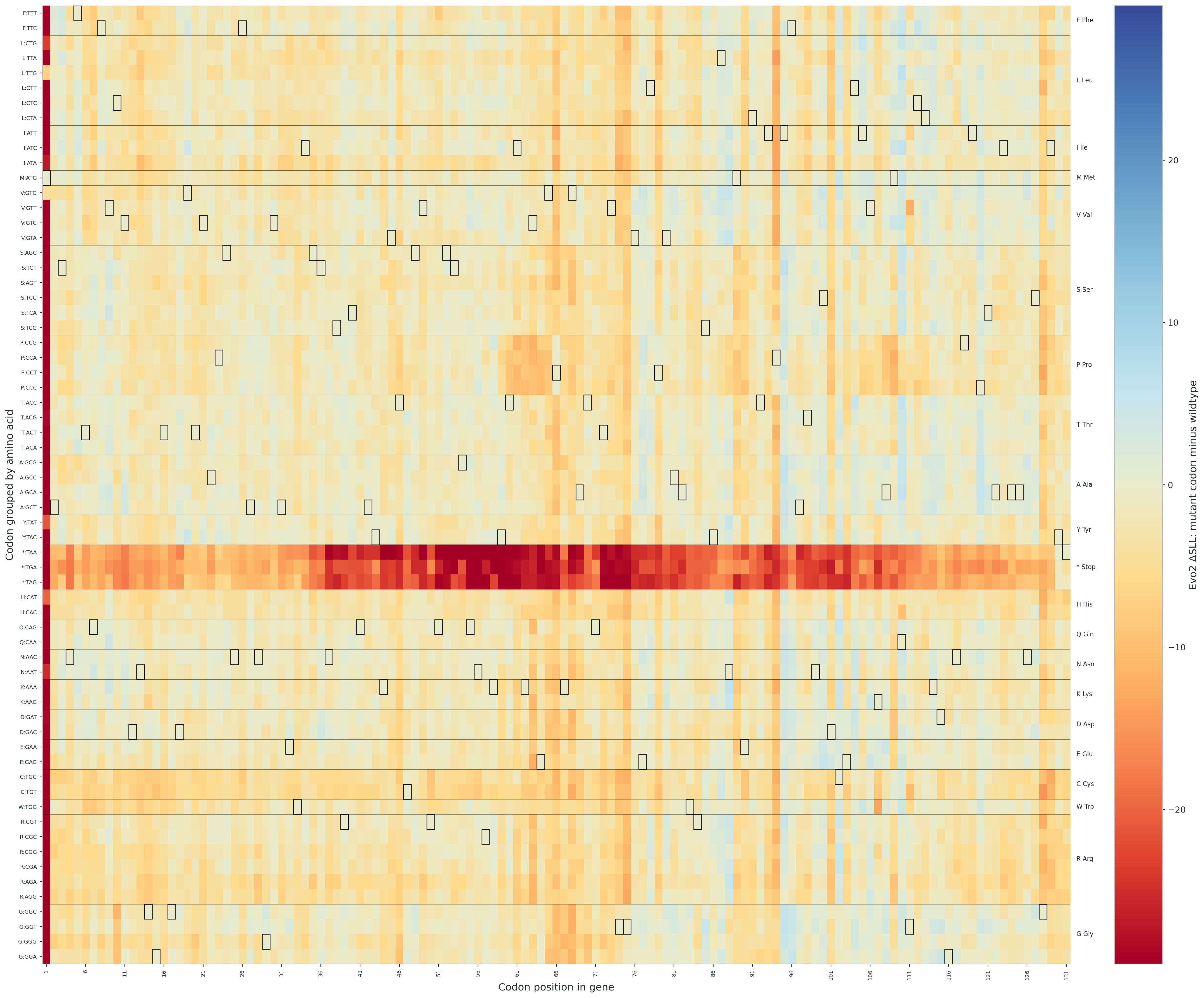

**Figure S2B. Codon-resolution heatmaps of Evo2** **ΔSLL scores across the MS2 coat gene.** The y-axis shows the substituted codon, grouped by encoded amino acid and labeled by nucleotide triplet, and the x-axis shows codon position within the indicated gene. Each cell reports the ΔSLL of the full mutant genome after replacement of the indicated codon at the indicated site. Rows within each amino acid block are ordered by *E. coli* codon usage, with the most frequently used codon for that amino acid at the top and descending frequencies beneath. Boxes outline wildtype codons. The heatmap provides codon-level resolution of the genome-wide scoring patterns shown in Fig. 1 and allow localized inspection of tolerated and strongly disfavored substitutions within each gene.

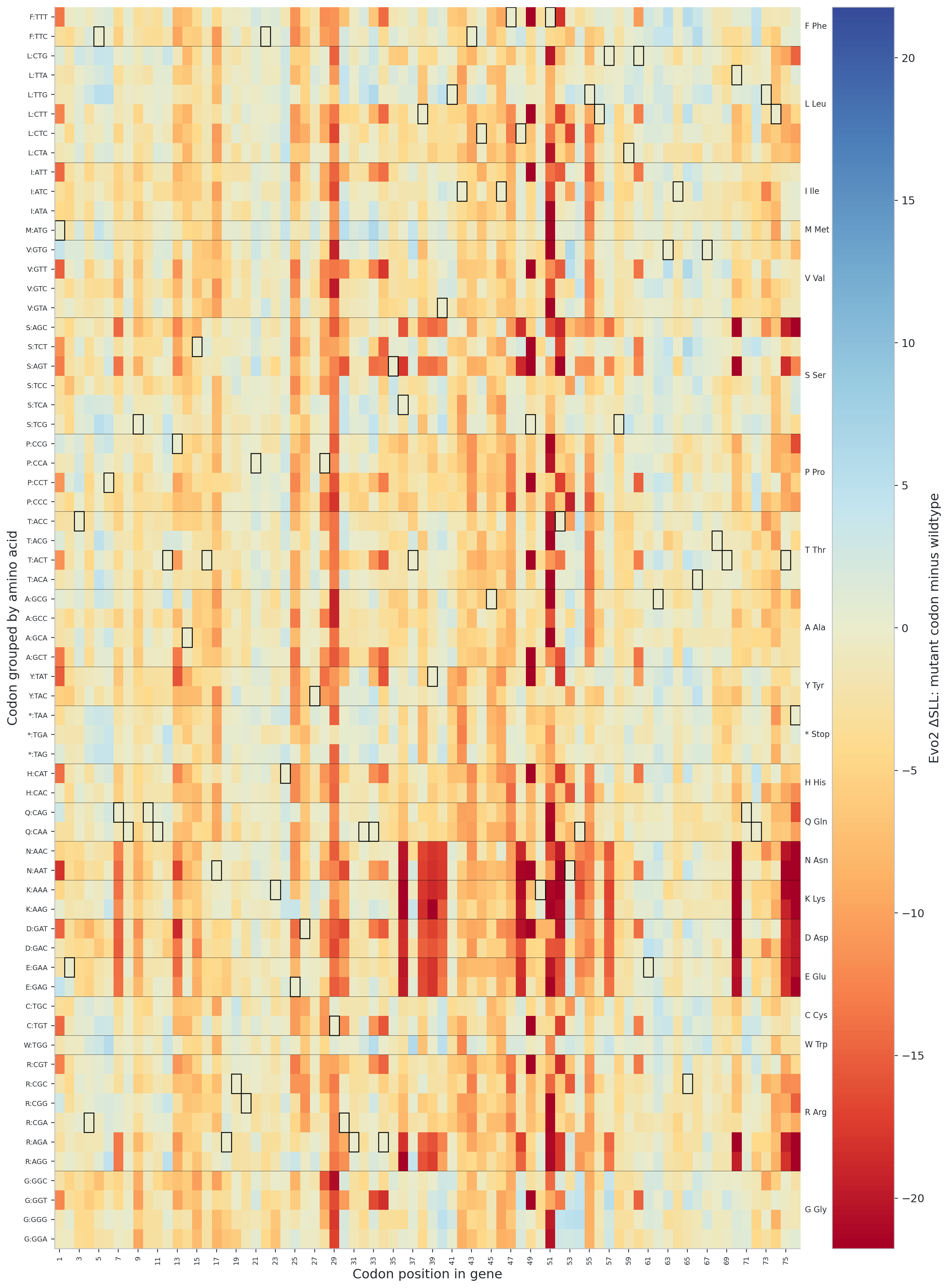

**Figure S2C. Codon-resolution heatmaps of Evo2 ΔSLL scores across the MS2 lysis gene.** The y-axis shows the substituted codon, grouped by encoded amino acid and labeled by nucleotide triplet, and the x-axis shows codon position within the indicated gene. Each cell reports the ΔSLL of the full mutant genome after replacement of the indicated codon at the indicated site. The heatmap provides codon-level resolution of the genome-wide scoring patterns shown in Fig. 1 and allows localized inspection of tolerated and strongly disfavored substitutions within each gene.

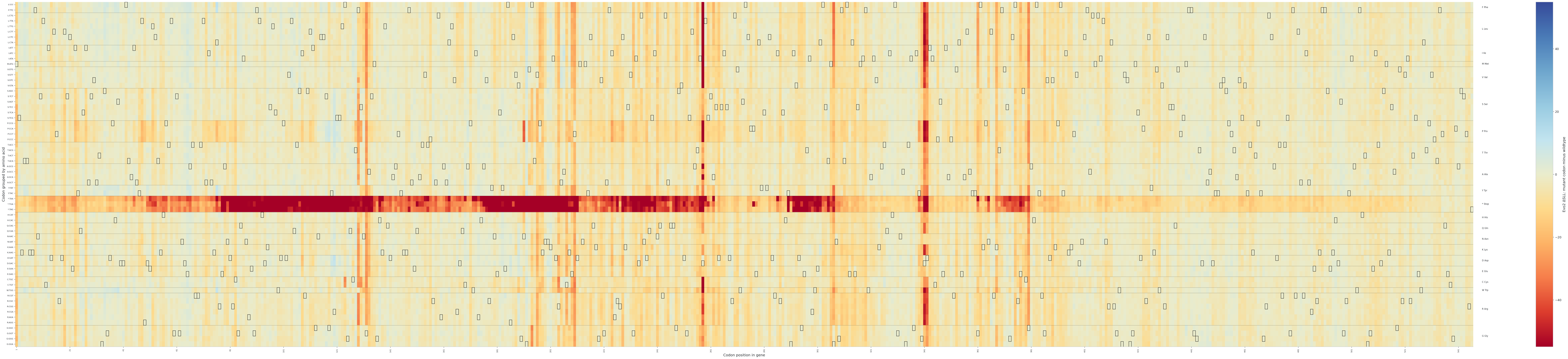

**Figure S2D. Codon-resolution heatmaps of Evo2** **ΔSLL scores across the MS2 replicase gene.** The y-axis shows the substituted codon, grouped by encoded amino acid and labeled by nucleotide triplet, and the x-axis shows codon position within the indicated gene. Each cell reports the ΔSLL of the full mutant genome after replacement of the indicated codon at the indicated site. Rows within each amino acid block are ordered by *E. coli* codon usage, with the most frequently used codon for that amino acid at the top and descending frequencies beneath. Boxes outline wildtype codons. The heatmap provides codon-level resolution of the genome-wide scoring patterns shown in Fig. 1 and allow localized inspection of tolerated and strongly disfavored substitutions within each gene.

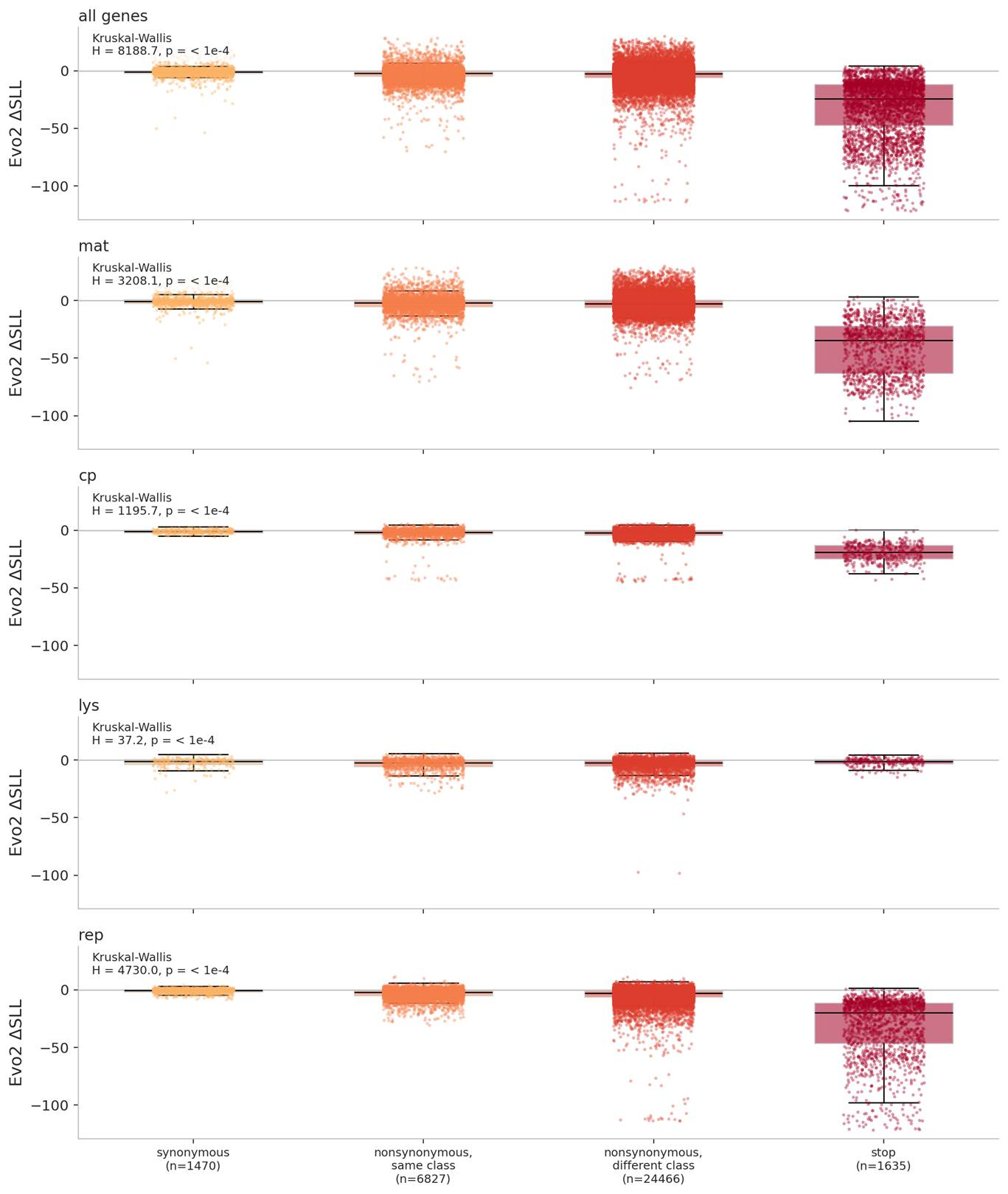

**Figure S3. Distribution of Evo2 ΔSLL scores across mutation classes in MS2 genes.** In each panel, the y-axis shows ΔSLL, the difference between the sequence log-likelihood of the mutant genome and that of the reference genome, such that more negative values indicate variants that Evo2 considers less plausible than the reference. Mutation classes are shown in the same order in every panel: synonymous substitutions, nonsynonymous substitutions that preserve amino-acid class, nonsynonymous substitutions that cross amino-acid class, and nonsense substitutions introducing stop codons. Panels are shown for all genes combined and separately for the maturation, coat, lysis, and replicase genes. The Kruskal–Wallis statistic shown in each panel asks whether the four mutation classes differ overall in their ΔSLL distributions. These plots show that Evo2 broadly distinguishes mutation classes across MS2, while also highlighting gene-specific departures from that pattern, particularly in the lysis gene.

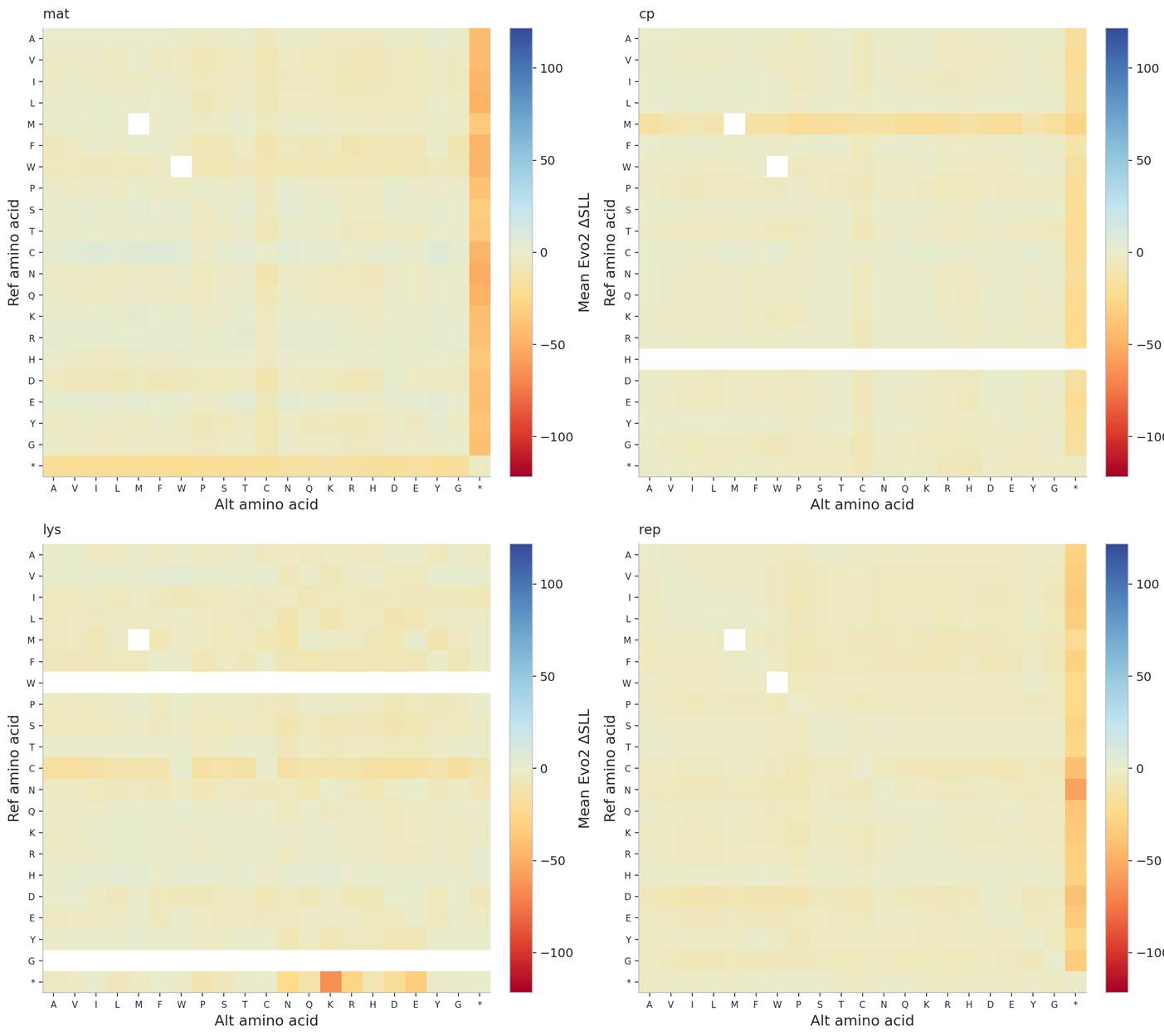

**Figure S4. Amino-acid substitution heatmaps derived from genome-wide Evo2 scoring of MS2 codon variants.** Each panel summarizes Evo2 ΔSLL scores for one MS2 gene by aggregating codon substitutions at the amino-acid level. The y-axis gives the reference amino acid present in the gene, and the x-axis gives the alternative amino acid encoded after mutation. Each cell therefore represents the mean ΔSLL across all codon substitutions in that gene that convert the indicated reference amino acid to the indicated alternative amino acid. More negative values (red) indicate substitutions that Evo2 considers less plausible relative to the reference genome, whereas more positive values (blue) indicate substitutions scored as closer to or more plausible than the reference. In practical terms, a red cell can be read as a more strongly penalized class of amino-acid replacement in that gene. Blank or white cells indicate that no substitutions of the corresponding reference-to-alternative amino-acid class were represented in that gene. This can occur because the gene does not contain the indicated reference amino acid, because no codon change in the enumerated set produces that amino-acid transition, or because the transition is undefined (for example, methionine-to-methionine under the standard genetic code, which has no synonymous codon alternative). The panels therefore provide a compressed view of gene-specific substitution preferences but should be interpreted alongside the codon-level plots, since each cell averages over potentially many distinct nucleotide changes and positions.

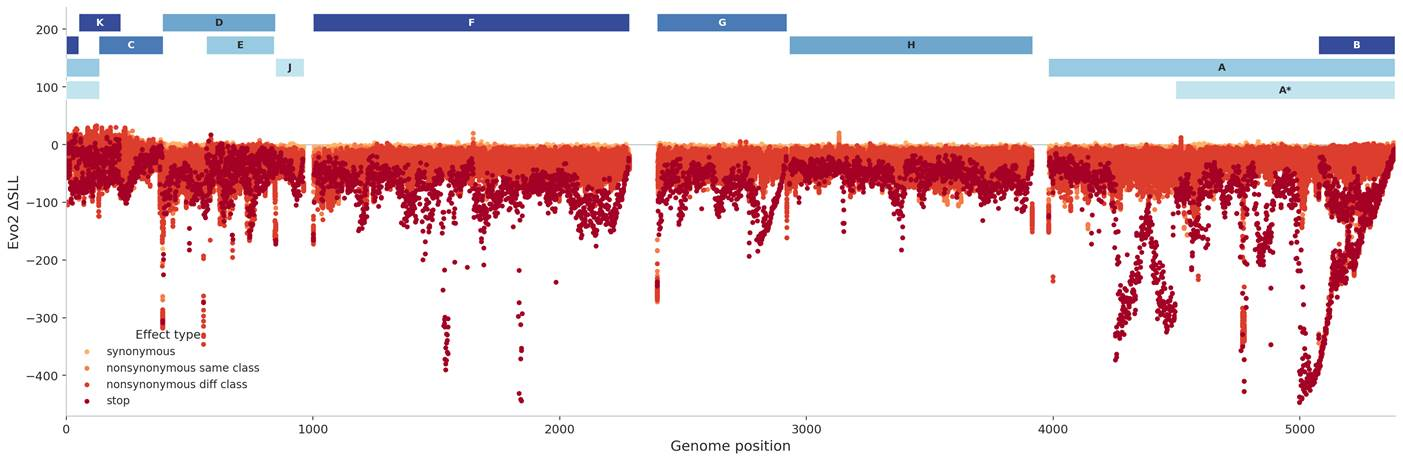

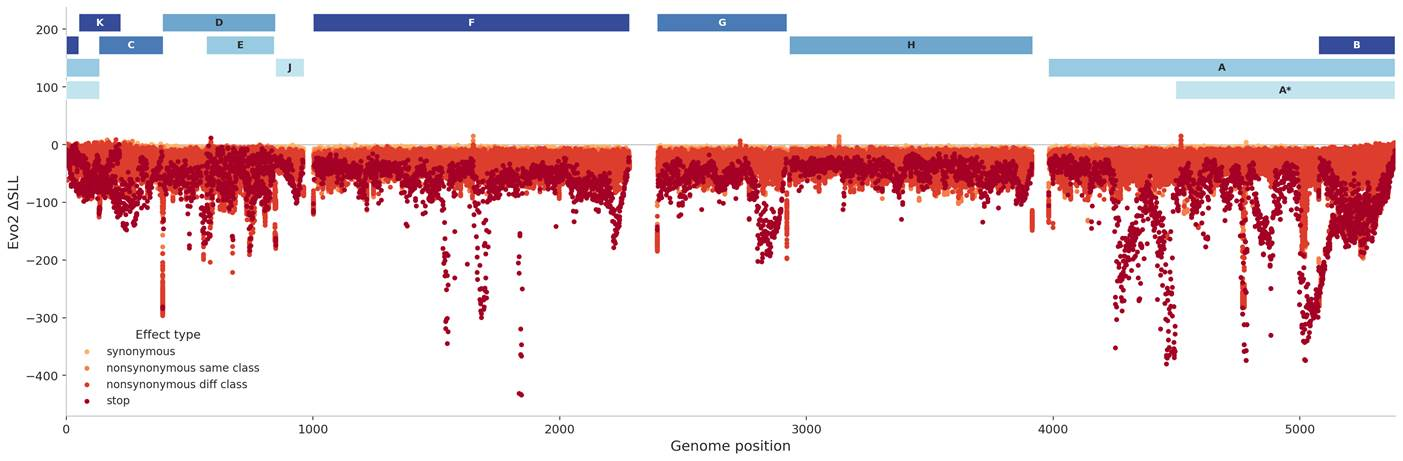

**Figure S5. Forward and reverse-complement Evo2 scoring for genome-wide** **ΦX174 codon substitutions.** Top, standard forward-pass ΔSLL analysis. Bottom, reverse-complement-averaged ΔSLL analysis using Evo2’s average_reverse_complement option. As in MS2, reverse-complement averaging reduces score variability at the 5′ end that arises from limited upstream context in autoregressive scoring. In ΦX174, however, this positional artifact is less visually disruptive than in MS2 and does not change the overall interpretation of the genome-wide scoring patterns.

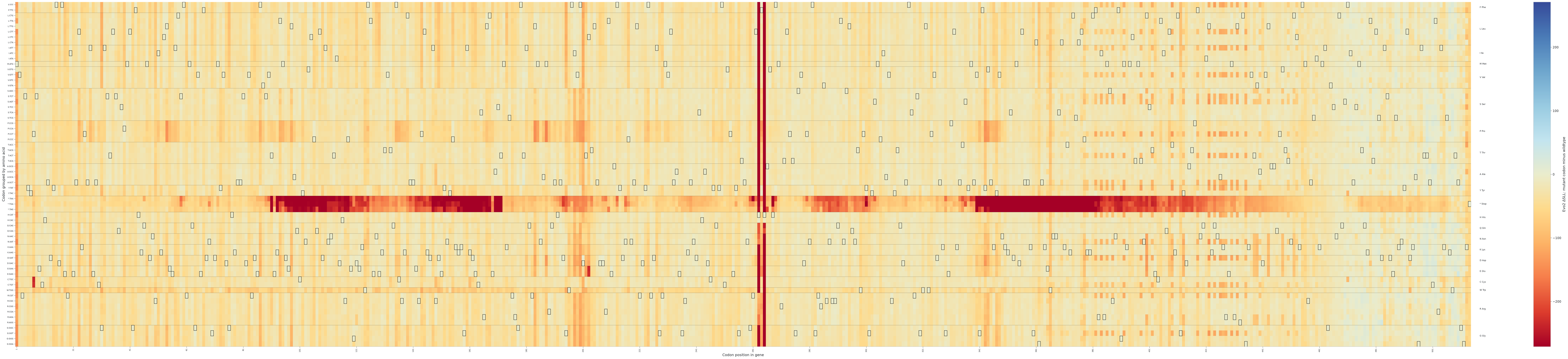

**Figure S6A. Codon-resolution heatmap of Evo2** **ΔSLL scores across the ΦX174 *A* gene (replication initiator and terminator).** The y-axis shows the substituted codon, grouped by encoded amino acid and labeled by nucleotide triplet, and the x-axis shows codon position within the indicated gene. Each cell reports the ΔSLL of the full mutant genome after replacement of the indicated codon at the indicated site. Rows within each amino acid block are ordered by *E. coli* codon usage, with the most frequently used codon for that amino acid at the top and descending frequencies beneath. Boxes outline wildtype codons. The heatmap provides codon-level resolution of the genome-wide scoring patterns shown in Fig. 4 and allows localized inspection of tolerated and strongly disfavored substitutions within the gene.

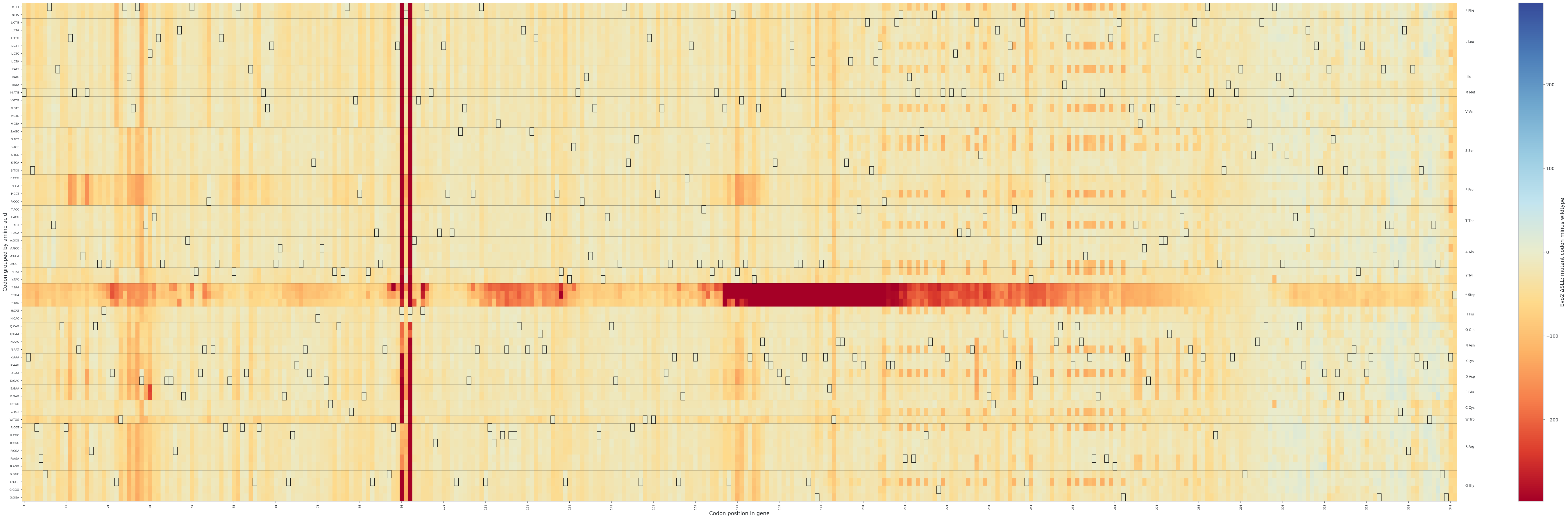

**Figure S6B. Codon-resolution heatmap of Evo2 ΔSLL scores across the** **ΦX174 *A** gene (host replication inhibitor).** The y-axis shows the substituted codon, grouped by encoded amino acid and labeled by nucleotide triplet, and the x-axis shows codon position within the indicated gene. Each cell reports the ΔSLL of the full mutant genome after replacement of the indicated codon at the indicated site. Rows within each amino acid block are ordered by *E. coli* codon usage, with the most frequently used codon for that amino acid at the top and descending frequencies beneath. Boxes outline wildtype codons. The heatmap provides codon-level resolution of the genome-wide scoring patterns shown in Fig. 4 and allows localized inspection of tolerated and strongly disfavored substitutions within the gene.

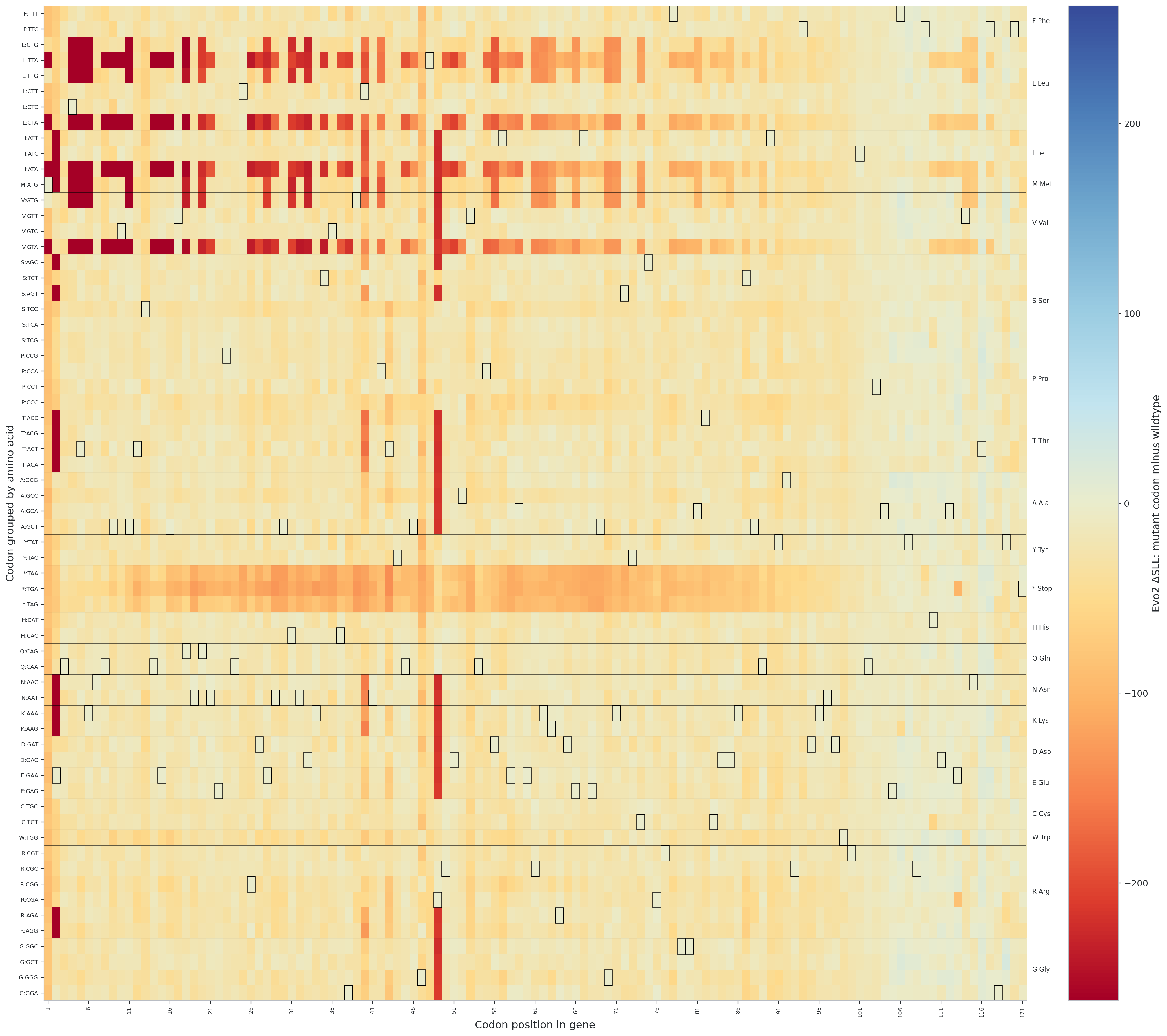

**Figure S6C. Codon-resolution heatmap of Evo2 ΔSLL scores across the ΦX174 *B* gene (minor internal scaffolding protein).** The y-axis shows the substituted codon, grouped by encoded amino acid and labeled by nucleotide triplet, and the x-axis shows codon position within the indicated gene. Each cell reports the ΔSLL of the full mutant genome after replacement of the indicated codon at the indicated site. Rows within each amino acid block are ordered by *E. coli* codon usage, with the most frequently used codon for that amino acid at the top and descending frequencies beneath. Boxes outline wildtype codons. The heatmap provides codon-level resolution of the genome-wide scoring patterns shown in Fig. 4 and allows localized inspection of tolerated and strongly disfavored substitutions within the gene.

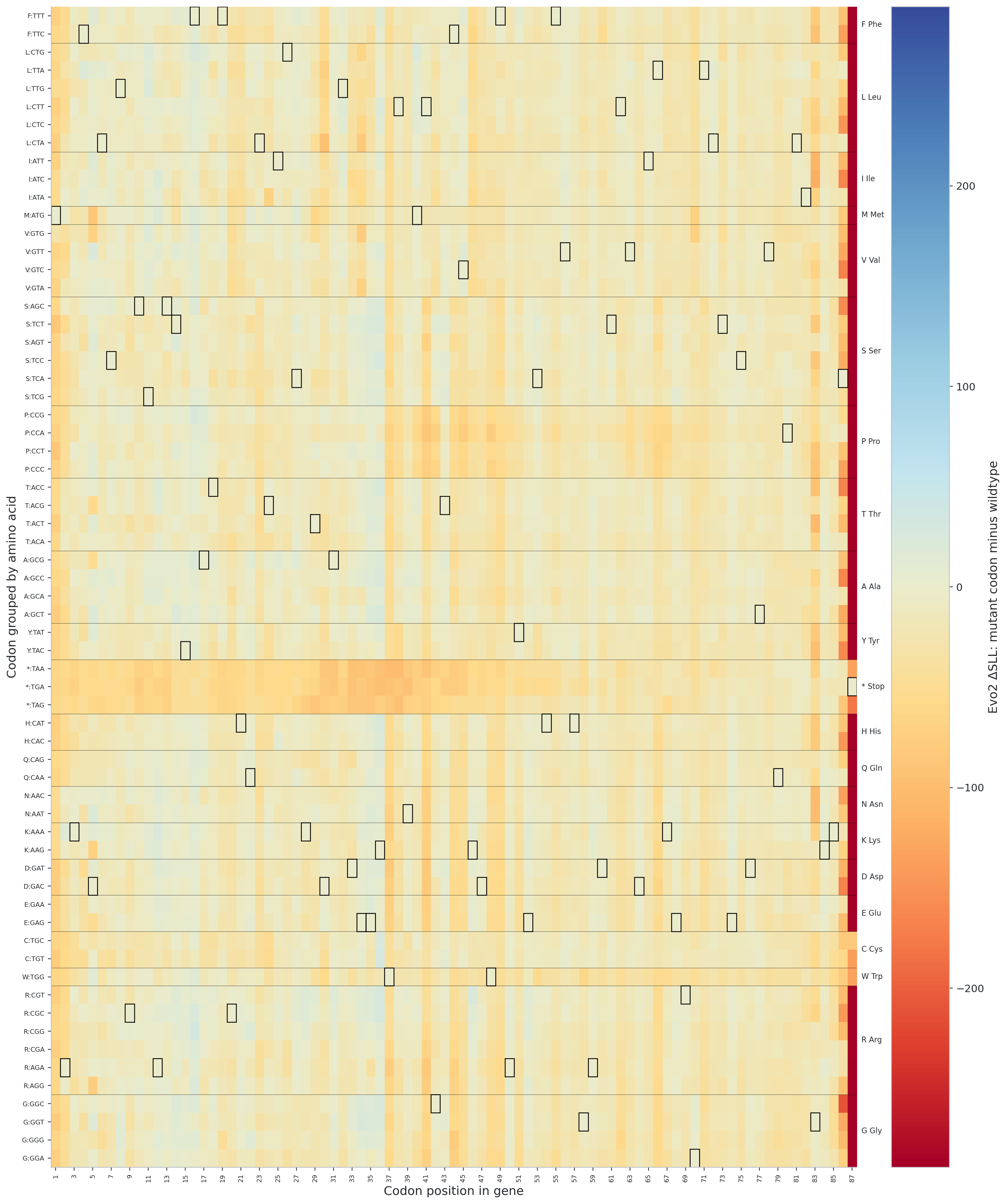

**Figure S6D. Codon-resolution heatmap of Evo2 ΔSLL scores across the ΦX174 *C* gene (replication regulator).** The y-axis shows the substituted codon, grouped by encoded amino acid and labeled by nucleotide triplet, and the x-axis shows codon position within the indicated gene. Each cell reports the ΔSLL of the full mutant genome after replacement of the indicated codon at the indicated site. Rows within each amino acid block are ordered by *E. coli* codon usage, with the most frequently used codon for that amino acid at the top and descending frequencies beneath. Boxes outline wildtype codons. The heatmap provides codon-level resolution of the genome-wide scoring patterns shown in Fig. 4 and allows localized inspection of tolerated and strongly disfavored substitutions within the gene.

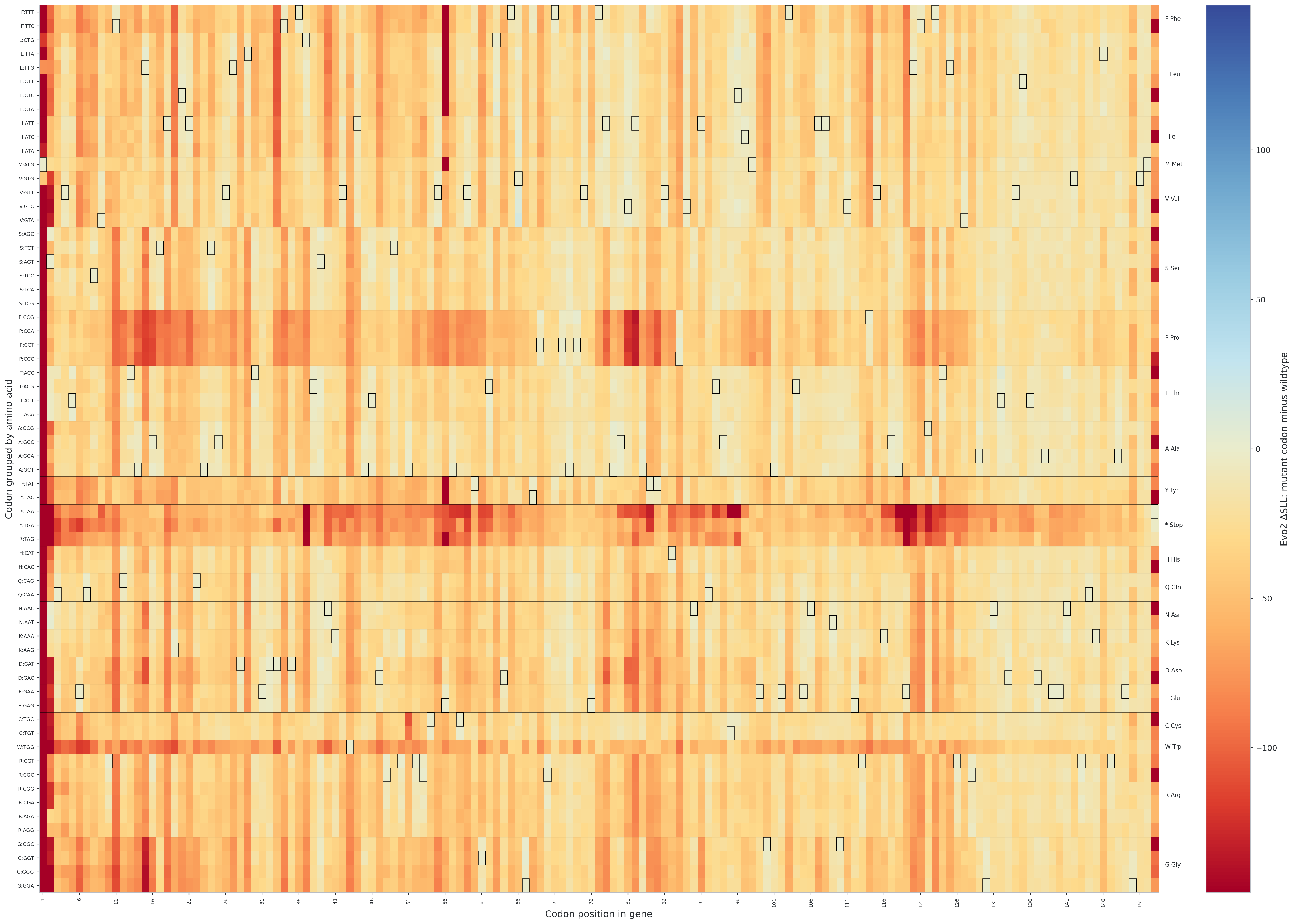

**Figure S6E. Codon-resolution heatmap of Evo2 ΔSLL scores across the ΦX174 *D* gene (DNA packaging protein).** The y-axis shows the substituted codon, grouped by encoded amino acid and labeled by nucleotide triplet, and the x-axis shows codon position within the indicated gene. Each cell reports the ΔSLL of the full mutant genome after replacement of the indicated codon at the indicated site. Rows within each amino acid block are ordered by *E. coli* codon usage, with the most frequently used codon for that amino acid at the top and descending frequencies beneath. Boxes outline wildtype codons. The heatmap provides codon-level resolution of the genome-wide scoring patterns shown in Fig. 4 and allows localized inspection of tolerated and strongly disfavored substitutions within the gene.

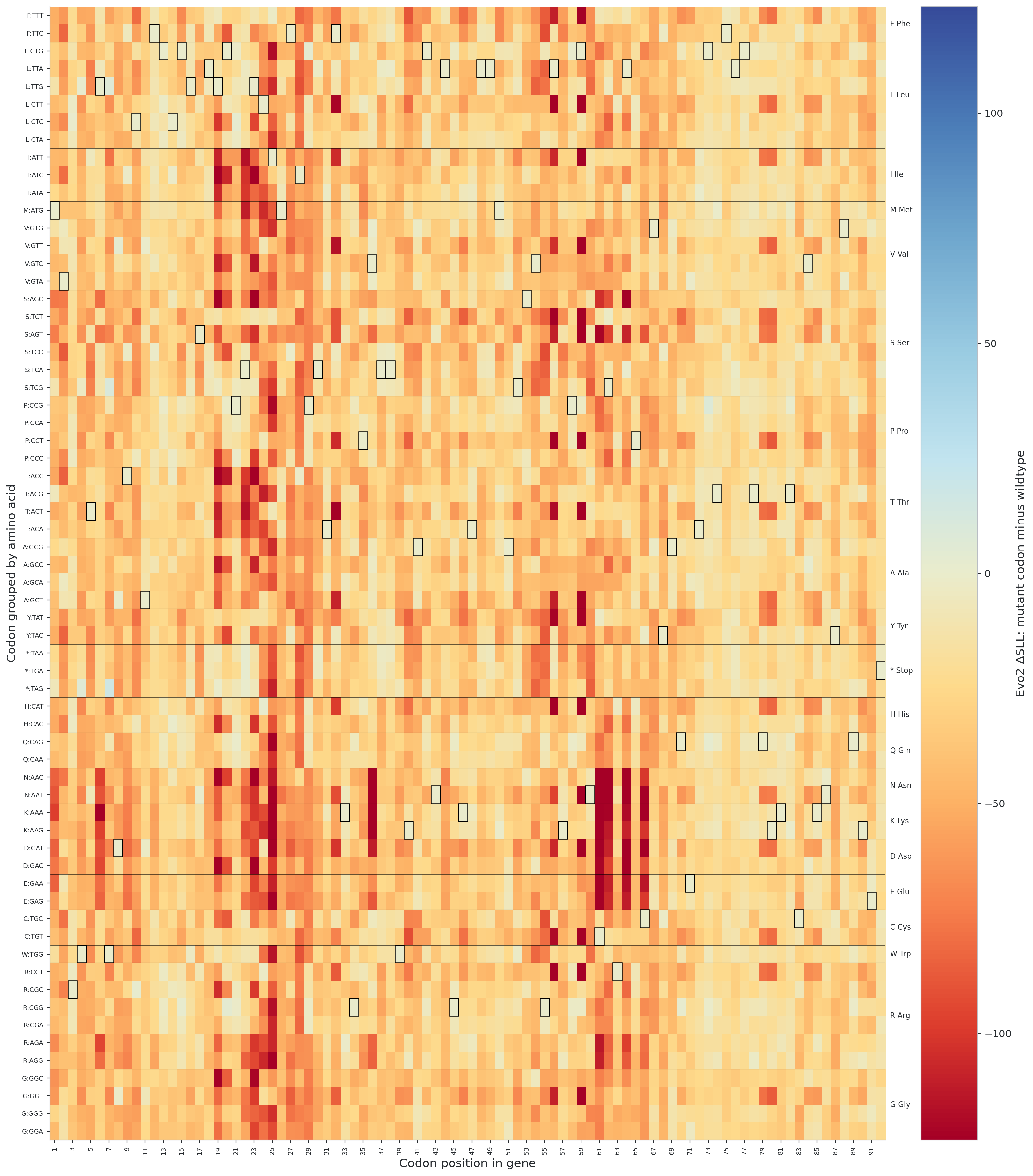

**Figure S6F. Codon-resolution heatmap of Evo2 ΔSLL scores across the ΦX174 *E* gene (lysis protein).** The y-axis shows the substituted codon, grouped by encoded amino acid and labeled by nucleotide triplet, and the x-axis shows codon position within the indicated gene. Each cell reports the ΔSLL of the full mutant genome after replacement of the indicated codon at the indicated site. Rows within each amino acid block are ordered by *E. coli* codon usage, with the most frequently used codon for that amino acid at the top and descending frequencies beneath. Boxes outline wildtype codons. The heatmap provides codon-level resolution of the genome-wide scoring patterns shown in Fig. 4 and allows localized inspection of tolerated and strongly disfavored substitutions within the gene.

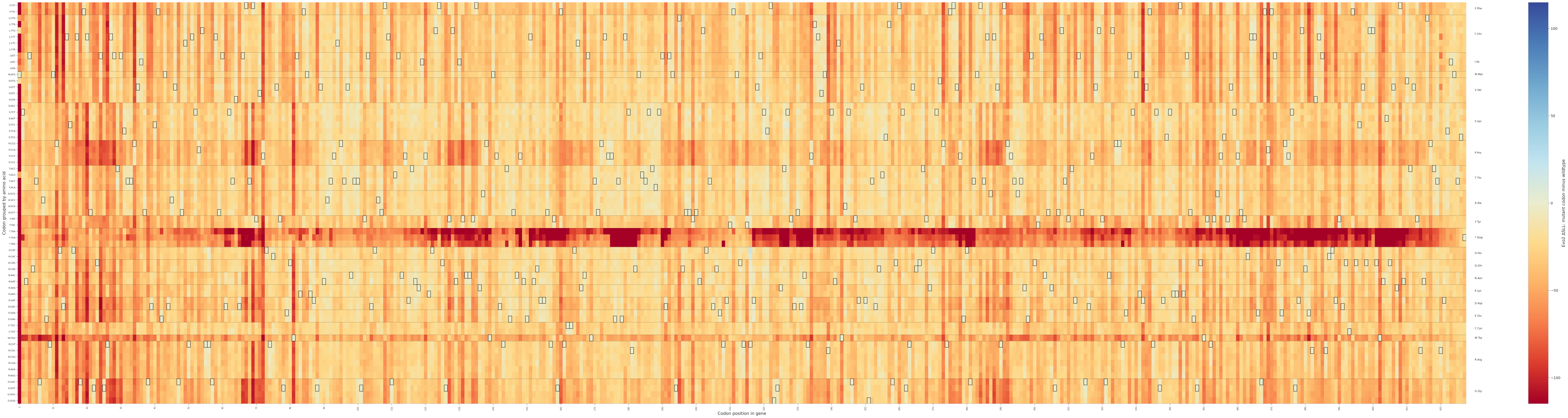

**Figure S6G. Codon-resolution heatmap of Evo2 ΔSLL scores across the ΦX174 *F* gene (major capsid protein).** The y-axis shows the substituted codon, grouped by encoded amino acid and labeled by nucleotide triplet, and the x-axis shows codon position within the indicated gene. Each cell reports the ΔSLL of the full mutant genome after replacement of the indicated codon at the indicated site. Rows within each amino acid block are ordered by *E. coli* codon usage, with the most frequently used codon for that amino acid at the top and descending frequencies beneath. Boxes outline wildtype codons. The heatmap provides codon-level resolution of the genome-wide scoring patterns shown in Fig. 4 and allows localized inspection of tolerated and strongly disfavored substitutions within the gene.

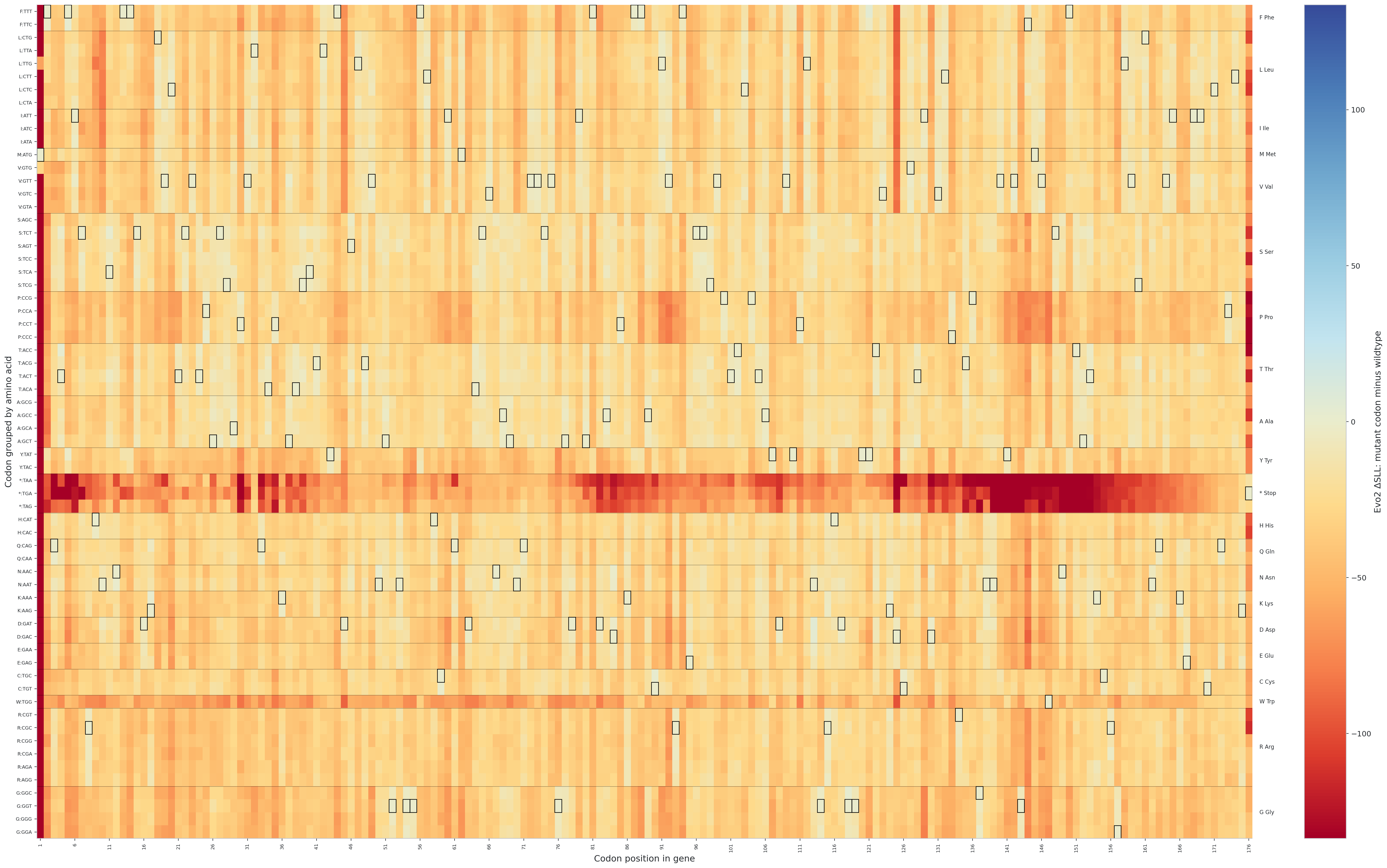

**Figure S6H. Codon-resolution heatmap of Evo2 ΔSLL scores across the ΦX174 *G* gene (major spike protein).** The y-axis shows the substituted codon, grouped by encoded amino acid and labeled by nucleotide triplet, and the x-axis shows codon position within the indicated gene. Each cell reports the ΔSLL of the full mutant genome after replacement of the indicated codon at the indicated site. Rows within each amino acid block are ordered by *E. coli* codon usage, with the most frequently used codon for that amino acid at the top and descending frequencies beneath. Boxes outline wildtype codons. The heatmap provides codon-level resolution of the genome-wide scoring patterns shown in Fig. 4 and allows localized inspection of tolerated and strongly disfavored substitutions within the gene.

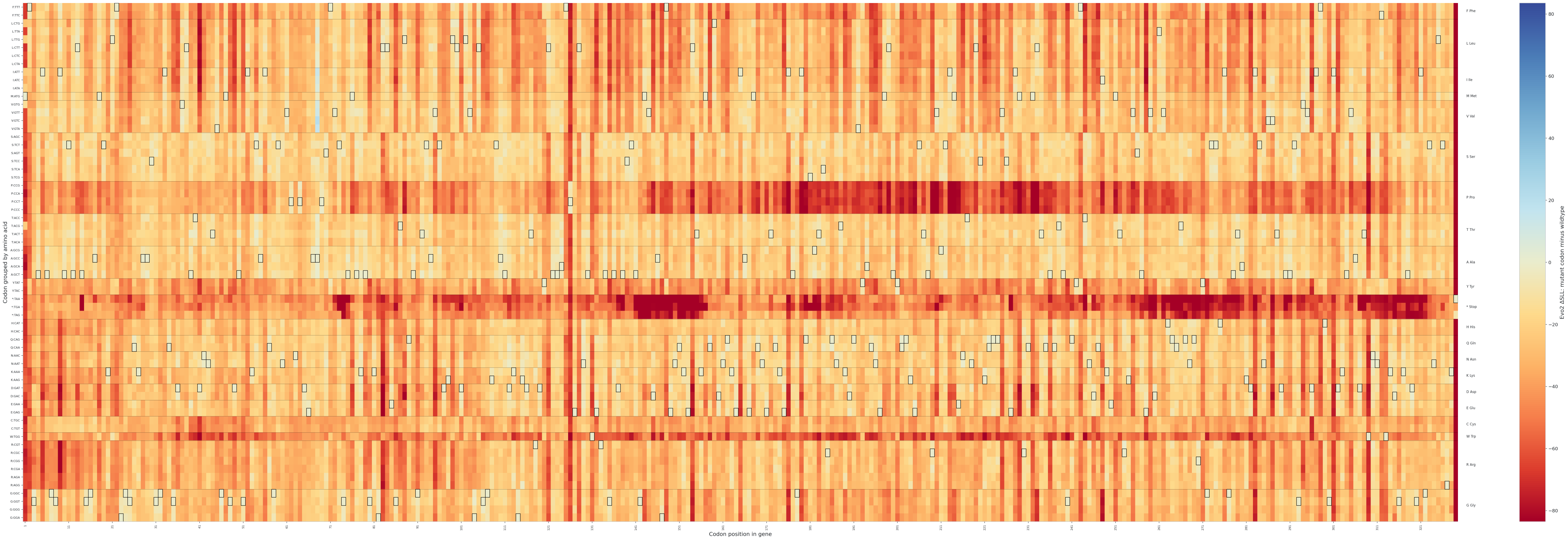

**Figure S6I. Codon-resolution heatmap of Evo2 ΔSLL scores across the ΦX174 *H* gene (DNA pilot protein).** The y-axis shows the substituted codon, grouped by encoded amino acid and labeled by nucleotide triplet, and the x-axis shows codon position within the indicated gene. Each cell reports the ΔSLL of the full mutant genome after replacement of the indicated codon at the indicated site. Rows within each amino acid block are ordered by *E. coli* codon usage, with the most frequently used codon for that amino acid at the top and descending frequencies beneath. Boxes outline wildtype codons. The heatmap provides codon-level resolution of the genome-wide scoring patterns shown in Fig. 4 and allows localized inspection of tolerated and strongly disfavored substitutions within the gene.

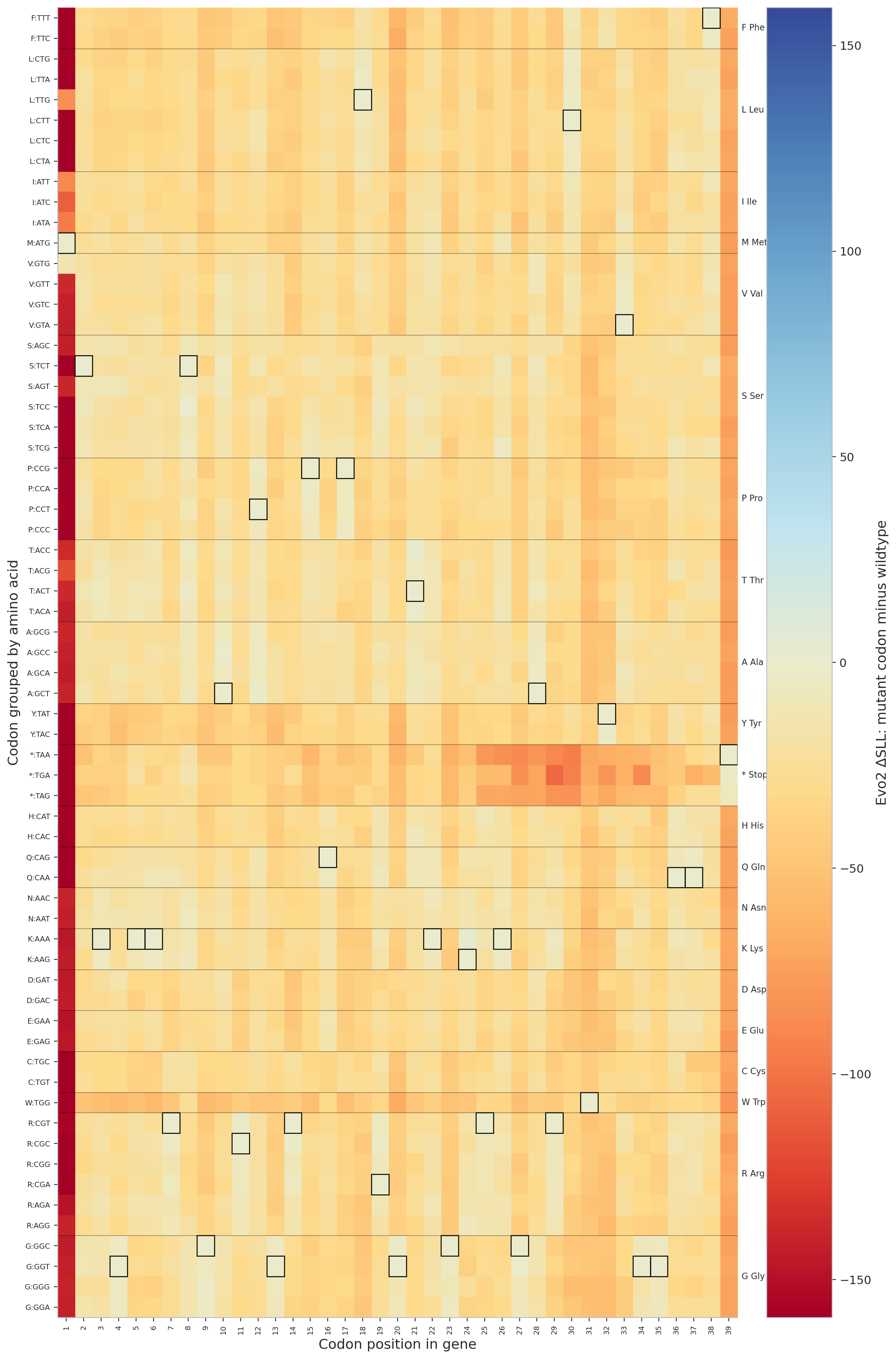

**Figure S6J. Codon-resolution heatmap of Evo2 ΔSLL scores across the ΦX174 *J* gene (DNA packaging / host-interaction protein).** The y-axis shows the substituted codon, grouped by encoded amino acid and labeled by nucleotide triplet, and the x-axis shows codon position within the indicated gene. Each cell reports the ΔSLL of the full mutant genome after replacement of the indicated codon at the indicated site. Rows within each amino acid block are ordered by *E. coli* codon usage, with the most frequently used codon for that amino acid at the top and descending frequencies beneath. Boxes outline wildtype codons. The heatmap provides codon-level resolution of the genome-wide scoring patterns shown in Fig. 4 and allows localized inspection of tolerated and strongly disfavored substitutions within the gene.

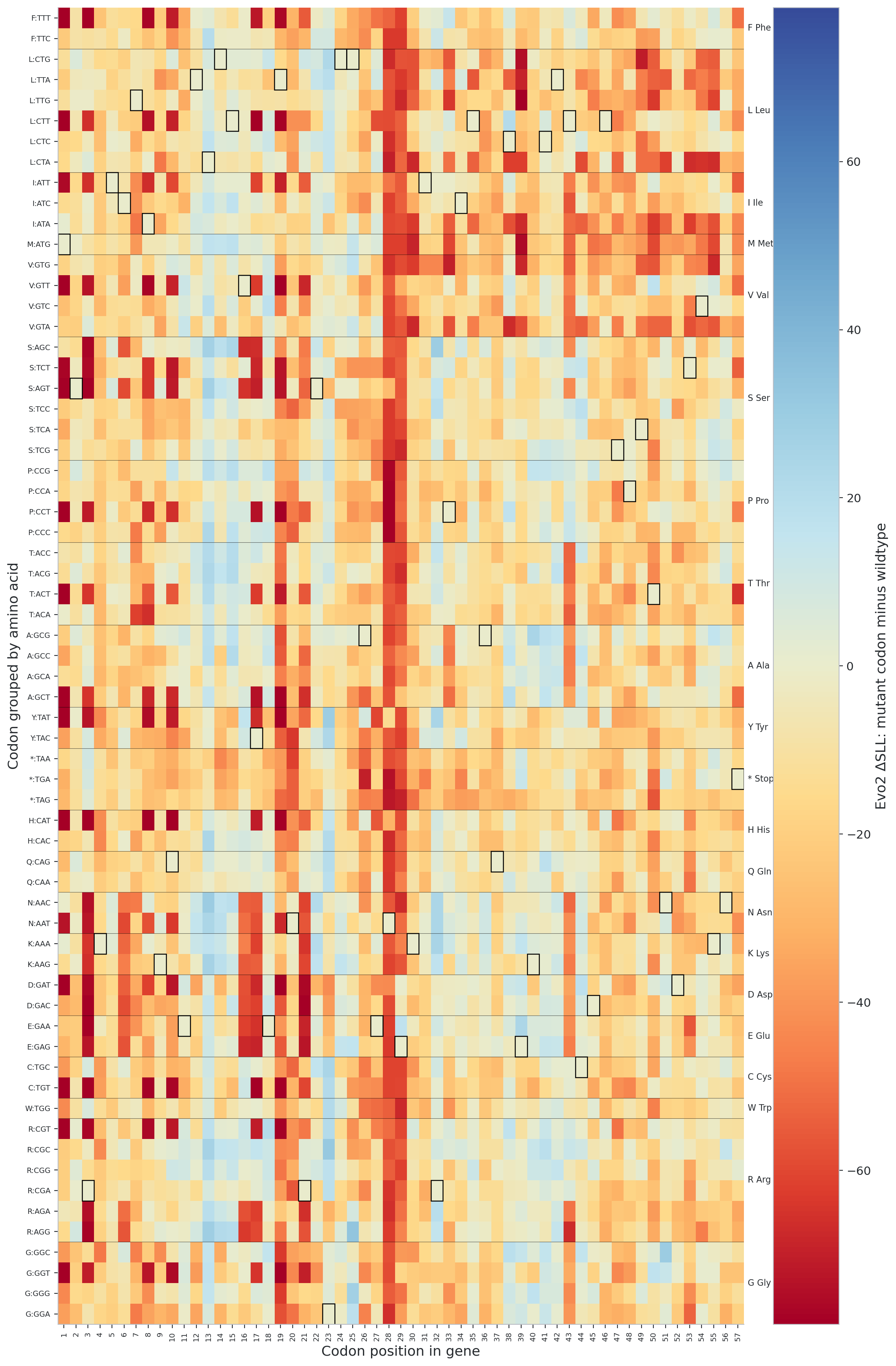

**Figure S6J. Codon-resolution heatmap of Evo2 ΔSLL scores across the ΦX174 *K* gene (small internal gene embedded within *A* and *C*).** The y-axis shows the substituted codon, grouped by encoded amino acid and labeled by nucleotide triplet, and the x-axis shows codon position within the indicated gene. Each cell reports the ΔSLL of the full mutant genome after replacement of the indicated codon at the indicated site. Rows within each amino acid block are ordered by *E. coli* codon usage, with the most frequently used codon for that amino acid at the top and descending frequencies beneath. Boxes outline wildtype codons. The heatmap provides codon-level resolution of the genome-wide scoring patterns shown in Fig. 4 and allows localized inspection of tolerated and strongly disfavored substitutions within the gene.

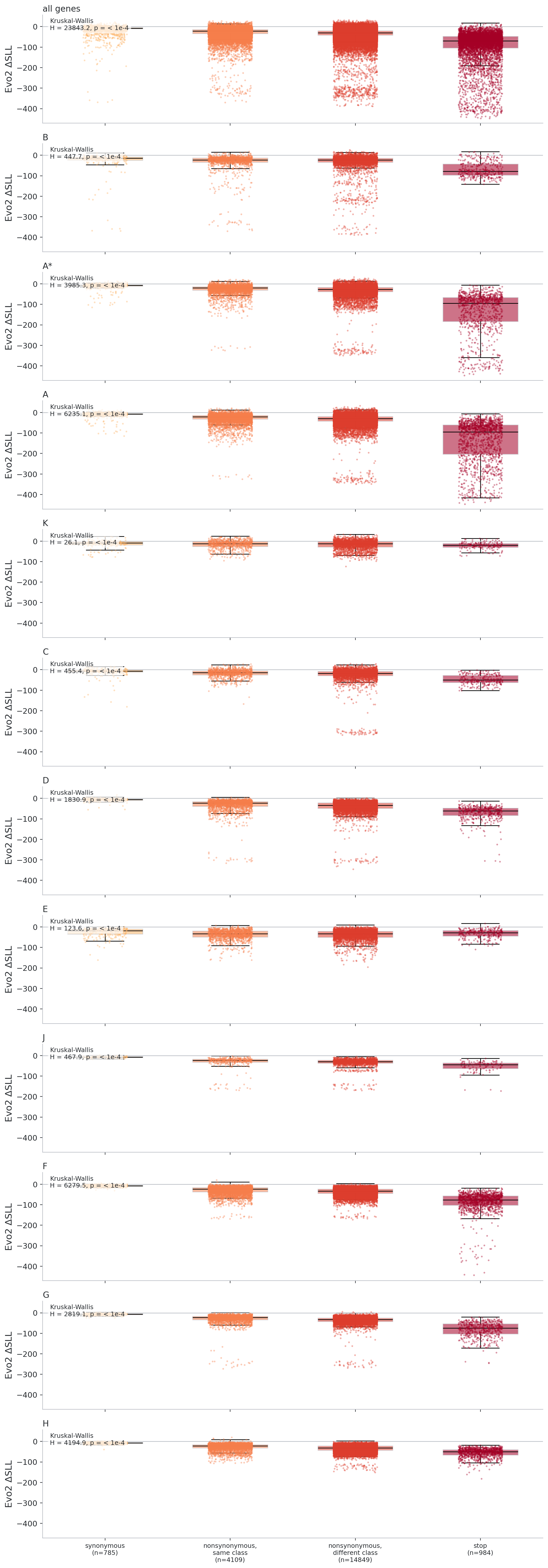

**Figure S7. Distribution of Evo2 ΔSLL scores across mutation classes in** **ΦX174 genes.** In each panel, the y-axis shows ΔSLL, the difference between the sequence log-likelihood of the mutant genome and that of the reference genome, such that more negative values indicate variants that Evo2 considers less plausible than the reference. Mutation classes are shown in the same order in every panel: synonymous substitutions, nonsynonymous substitutions that preserve amino-acid class, nonsynonymous substitutions that cross amino-acid class, and nonsense substitutions introducing stop codons. Panels are shown for genes are in the same order as in Fig. S6. The Kruskal–Wallis statistic shown in each panel asks whether the four mutation classes differ overall in their ΔSLL distributions.

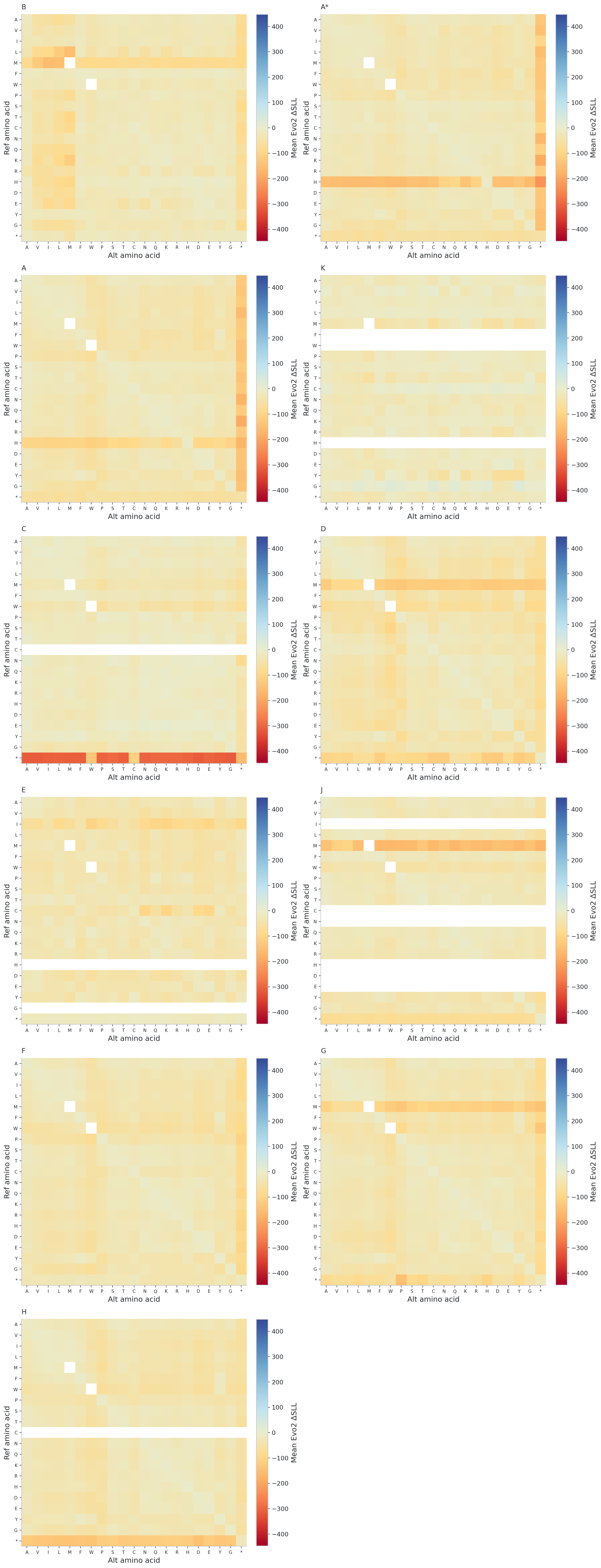

**Figure S8. Amino-acid substitution heatmaps derived from genome-wide Evo2 scoring of ΦX174** **codon variants.** Each panel summarizes Evo2 ΔSLL scores for one ΦX174 gene by aggregating codon substitutions at the amino-acid level. The y-axis gives the reference amino acid present in the gene, and the x-axis gives the alternative amino acid encoded after mutation. Each cell therefore represents the mean ΔSLL across all codon substitutions in that gene that convert the indicated reference amino acid to the indicated alternative amino acid. More negative values indicate substitutions that Evo2 considers less plausible relative to the reference genome, whereas more positive values indicate substitutions scored as closer to or more plausible than the reference. Blank cells indicate amino-acid transitions not represented in the corresponding gene. These panels provide an amino-acid-level summary of the codon-resolution patterns shown in Fig. S6 and allow comparison of broad substitution preferences across ΦX174 genes.

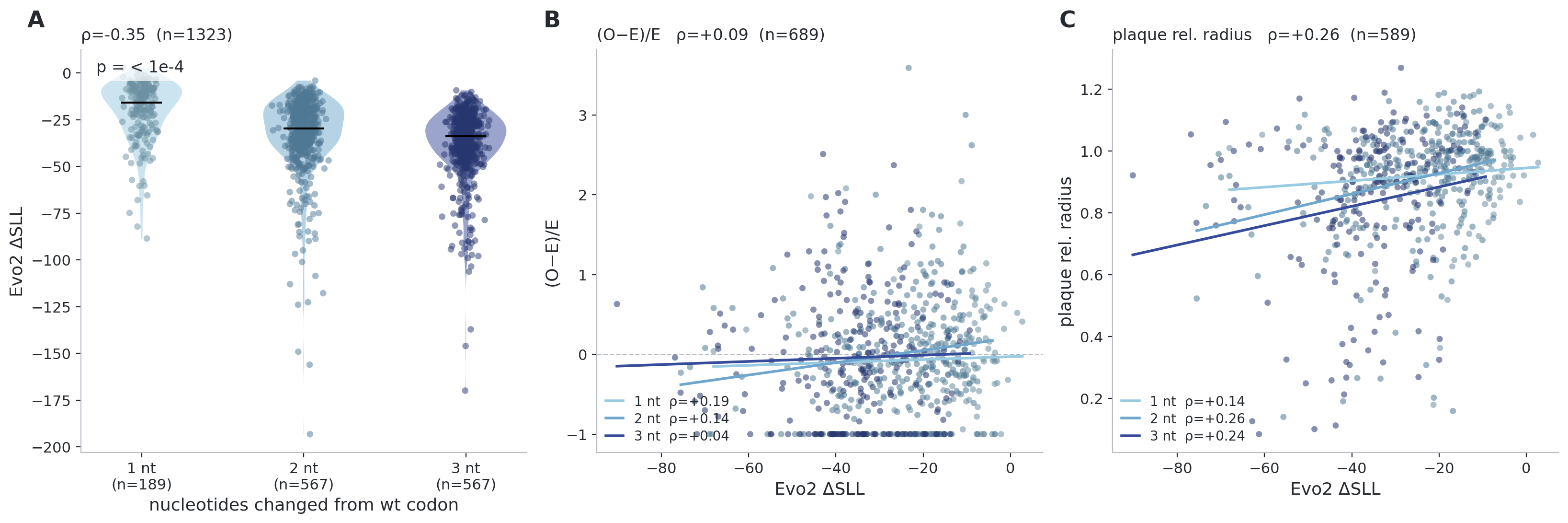

**Figure S9. Reference-distance effects in the** **ΦX174 gene *G* benchmark.** **(A)** ΔSLL values grouped by the number of nucleotide differences separating each mutant codon from the wild-type codon (1, 2, or 3). This panel shows that codons closer to the reference generally receive higher Evo2 plausibility scores. **(B)** Relationship between ΔSLL and the enrichment-like metric (observed - expected)/expected, shown separately for codons differing by one, two, or three nucleotides from the reference. **(C)** Relationship between  ΔSLL and relative plaque radius, shown separately for codons differing by one, two, or three nucleotides from the reference.
